# Global analysis of the yeast knock-out phenome

**DOI:** 10.1101/2022.12.22.521593

**Authors:** Gina Turco, Christie Chang, Rebecca Y. Wang, Griffin Kim, Emily Stoops, Brianna Richardson, Vanessa Sochat, Jennifer Rust, Rose Oughtred, Nathaniel Thayer, Fan Kang, Michael S. Livstone, Sven Heinicke, Mark Schroeder, Kara J. Dolinski, David Botstein, Anastasia Baryshnikova

**Affiliations:** Calico Life Sciences LLC; South San Francisco, CA, USA; Lewis-Sigler Institute for Integrative Genomics, Princeton University; Princeton, NJ, USA; Lawrence Livermore National Laboratory; Livermore, CA, USA

**Author notes:** Department of Computer & Information Science & Engineering, University of Florida; Gainesville, FL, USA.

## Abstract

Genome-wide phenotypic screens in the budding yeast *Saccharomyces cerevisiae* have produced the largest, richest and most systematic phenotypic description of any organism. Such an achievement was enabled by the development of highly scalable phenotypic assays and construction of the yeast knock-out (YKO) collection, comprising ~5,000 isogenic strains each deleted for exactly one open reading frame. Systematic screening of the YKO collection led to ~500 publications describing ~14,500 phenotypes capturing nearly every aspect of yeast biology. Yet, integrative analyses of this rich data source have been virtually impossible due to the lack of a central repository and consistent meta-data annotations. Here, we describe the aggregation, harmonization and analysis of all published phenotypic screens of the YKO collection, which we refer to as the Yeast Phenome (www.yeastphenome.org). To demonstrate the power of data integration and illustrate how much it facilitates the generation of testable hypotheses, we present three discoveries uniquely enabled by Yeast Phenome. First, we use the variation in the number of phenotypes per gene to identify tryptophan homeostasis as a central point of vulnerability to a wide range of chemical compounds, including FDA-approved drugs. Second, using phenotypic profiles as a tool for predicting gene function, we identify and validate the role of *YHR045W* as a novel regulator of ergosterol biosynthesis and DNA damage response, and *YGL117W* as a new member of the aromatic amino acid biosynthesis pathway. Finally, we describe a surprising exponential relationship between phenotypic similarity and intergenic distance in both yeast and human genomes. This relationship, which stretches as far as 380 kb in yeast and 100 Mb in humans, suggests that gene positions are optimized for function to a much greater extent than appreciated previously. Overall, we show that Yeast Phenome enables systematic enquiries into the nature of gene-gene and gene-phenotype relationships and is an important new resource for systems biology.

## INTRODUCTION

Connecting genotypes to phenotypes is essential for understanding the molecular architecture of complex traits and developing successful therapies against aging and disease. The assembly of large human cohorts, coupled with deep phenotyping and advanced computational analysis, is enabling great progress towards uncovering genome-wide phenotypic associations in natural human populations (*1*). However, inferring causal gene-trait relationships from these associations remains a challenge due to the complexity of human genetics, physiology, socio-economic structure and environmental exposures. An orthogonal approach to map genes to phenotypes has long been available through model organisms that allow systematic gene-by-gene perturbations in isogenic backgrounds and carefully controlled experimental environments.

The budding yeast *Saccharomyces cerevisiae* has pioneered the systematic phenotypic analysis of gene perturbations (*2*). In 2002, a consortium of laboratories released the yeast knock-out (YKO) collection, which provided a complete set of isogenic strains each deleted for exactly one open reading frame (ORF) (*3*). This collection, along with progress in automation and parallelization, enabled rapid, affordable and comprehensive loss-of-function screens that examined nearly every aspect of yeast biology that could be measured on a large scale. However, the results of these screens remained physically scattered and disorganized, thus preventing systematic analysis and integration. In the absence of a central repository and consistent metadata annotations, it has been impossible to know exactly which experiments have been done, how they compare to one another, and what information they contribute to our global understanding of yeast as a complex biological system. Here, we address this problem and describe Yeast Phenome, a data library that aggregates and annotates all published screens of the YKO collection. Currently, Yeast Phenome contains ~43 million causal gene-to-phenotype links, which represents the largest, richest and most systematic phenotypic description of any organism. To encourage exploration, download and analysis, we have made all data and metadata available at www.yeastphenome.org.

The aggregation and harmonization of YKO data in Yeast Phenome provides a unique dataset that enables discoveries that could not be made with any single experiment in isolation. To demonstrate its value, we provide several examples of Yeast Phenome data analysis and describe three key findings, ranging from gene-level to system-level observations. First, we analyze the variation in the number of phenotypes per gene and find that tryptophan biosynthesis is an exceptional metabolic pathway that is required for resistance to over 1,000 different chemical compounds. Second, we show that a multidimensional phenotypic profile, i.e. the set of all known phenotypes associated with a gene, is a strong predictor of gene function that complements and reinforces other genomic datasets. Using phenotypic profiles as predictive tools, we identify and validate the roles of two uncharacterized ORFs (*YHR045W* and *YGL117W*). Finally, we uncover an unexpected relationship between phenotypic profile similarity and intergenic distance, which potentially reflects the functional architecture of yeast and human genomes. Overall, we show that data curation is a powerful approach for generating new datasets and identifying general patterns that are not apparent on a smaller scale.

## RESULTS

### Building a data library of knock-out phenotypic screens

The YKO is a collection of ~5,000 yeast strains where every ORF is individually deleted and replaced by a selectable marker linked to an ORF-specific molecular barcode in a common genetic background (fig. S1A–B). An exhaustive survey of the literature (Materials & Methods) showed that, between November 2000 and May 2022, 366 research groups published 531 studies, each describing the systematic testing of at least 1,000 haploid or homozygous diploid YKO mutants for one or more phenotypes under one or more experimental conditions (Fig. 1A). To examine the wealth of information concealed in these data, we curated the 531 publications and assembled a comprehensive compendium of 14,495 knock-out screens (Fig. 1A). We developed standard vocabularies to annotate and cross-reference 6,731 phenotypes and 7,536 experimental environments associated with the screens, and devised a reproducible computational pipeline for extracting, formatting and normalizing data from each publication (fig. S1B; Materials & Methods). Through close collaboration with 149 yeast researchers, we recovered 413 novel screens (3% of the total) corresponding to extended and, typically, more quantitative versions of previously published screens (Materials & Methods).

**Fig. 1:**
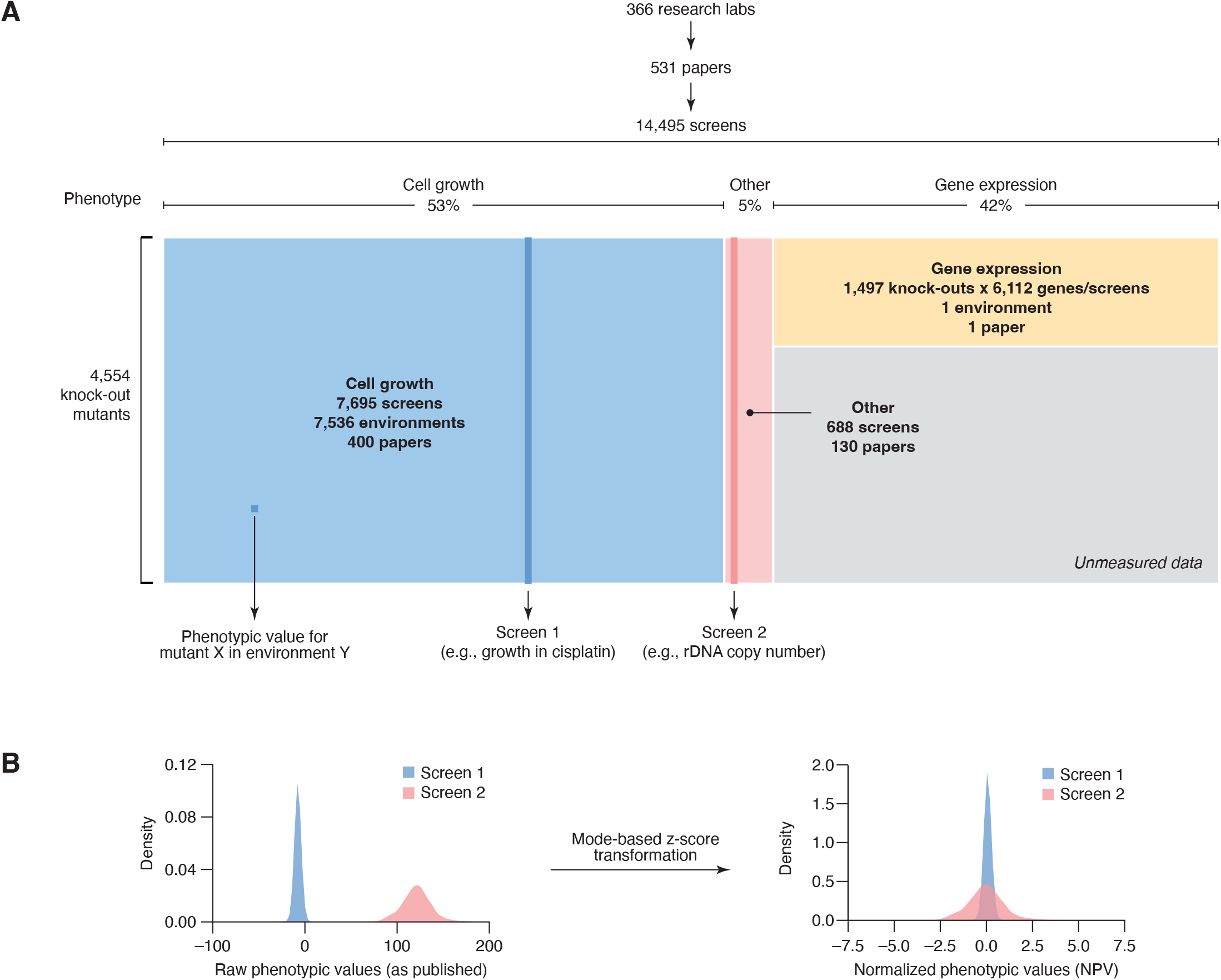
Yeast Phenome (www.yeastphenome.org) is a data library of published genomescale screens of the yeast knock-out (YKO) collection. (**A**) Yeast Phenome can be thought of as a data matrix where each row is a knock-out mutant and each column is a phenotypic screen. The matrix contains phenotypic values obtained by extracting data from 531 papers published by 366 research labs. The phenotypes tested by the screens and the experimental conditions/environments, in which the phenotypes were tested (e.g., chemical compounds, pH, temperature, growth media), were annotated using standard vocabularies. Three major classes of phenotypes (cell growth, gene expression and other) are highlighted in blue, yellow and pink, respectively. Grey represents unmeasured data because gene expression profiles were tested for only ~1,500 knock-out mutants. (**B**) To facilitate analysis and interpretation, raw phenotypic values (i.e., those released in the publication) were normalized using a modified z-score transformation which uses the mode (instead of the mean) and standard deviation from the mode to shift and scale the data.

The Yeast Phenome data library is experimentally and biologically diverse (Fig. 1A). The most tested phenotype (53% of all screens) is cell growth measured by colony size, optical density in liquid culture, relative barcode abundance in pools and several other metrics (Fig. 1A, blue). Because it is relatively easy to measure, growth of ~5,000 knock-out mutants has been tested in 7,536 different environments, most of which (96%) involved a chemical compound of known or unknown mode-of-action (Fig. 1A). For ~1,500 knock-out mutants (~30% of the YKO collection), growth measurements are supplemented by mRNA expression levels of 6,112 genes, representing the second most common phenotype in Yeast Phenome (42% of all screens; Fig. 1A, yellow). Despite being measured primarily in a single unperturbed environment (*4*), these genome-wide expression profiles of knock-out mutants provide a large and diverse set of molecular biomarkers that may act upstream of other phenotypes, including response to chemical treatments. The remaining 5% of screens in Yeast Phenome are a mosaic of 670 phenotypes that describe the state of the genome, proteome and metabolome of knock-out mutants, along with morphological parameters and other cellular phenotypes, such as protein localization and intracellular pH (Fig. 1A, red). These phenotypes are generally measured using advanced technologies (e.g., mass spectrometry, next-generation sequencing, high-resolution microscopy), complex reporter systems and, sometimes, longitudinal sampling, which probe yeast biology in greater detail but are limited in throughput (on average, 5.7 screens per publication). As such, they create many small but valuable datasets that are scattered throughout the literature and have never been examined in the context of other datasets. The inclusion of these data in Yeast Phenome creates the first opportunity to make novel insights based on their integration.

To facilitate the analysis and interpretation of diverse Yeast Phenome data, we implemented several conventions and normalizations (Materials & Methods). Most importantly, since different phenotypes followed dramatically different distributions but were consistently unimodal, we used the mode as a reference to normalize each screen using a modified z-score transformation (Fig. 1B, Materials & Methods). As a result, all phenotypic values reported in Yeast Phenome can be universally interpreted as standardized deviations from the most typical mutant, which, assuming extreme phenotypes are rare, is also likely to resemble the wild-type strain. Both original and transformed data, which we refer to as normalized phenotypic values (NPVs), are available at www.yeastphenome.org.

### Yeast Phenome data are reproducible and provide unique information about gene function

Thanks to its size and meta-data annotations, Yeast Phenome provides an opportunity to investigate the quality of YKO data and test their robustness to common sources of experimental noise. For example, we can easily identify and compare independent screens that examined the same phenotype under similar experimental conditions, and therefore assess the biological reproducibility of the phenotype. To demonstrate this point, we compared 8 independent screens of respiratory metabolism (i.e., growth on glycerol) and found that, on average, 71% of respiration-deficient mutants identified in any one screen were reproduced in at least 5 of the 8 replicates (note S1; fig. 2A). The similarity of the 8 screens was nearly complete (cosine ρ = 0.994 ± 0.003, mean ± std. dev.) when, instead of a gene-by-gene overlap, we compared the screens’ enrichment profiles across the genetic interaction similarity network using Spatial Analysis of Functional Enrichment (SAFE) (*5*) (note S1; fig. 2B).

**Fig. 2:**
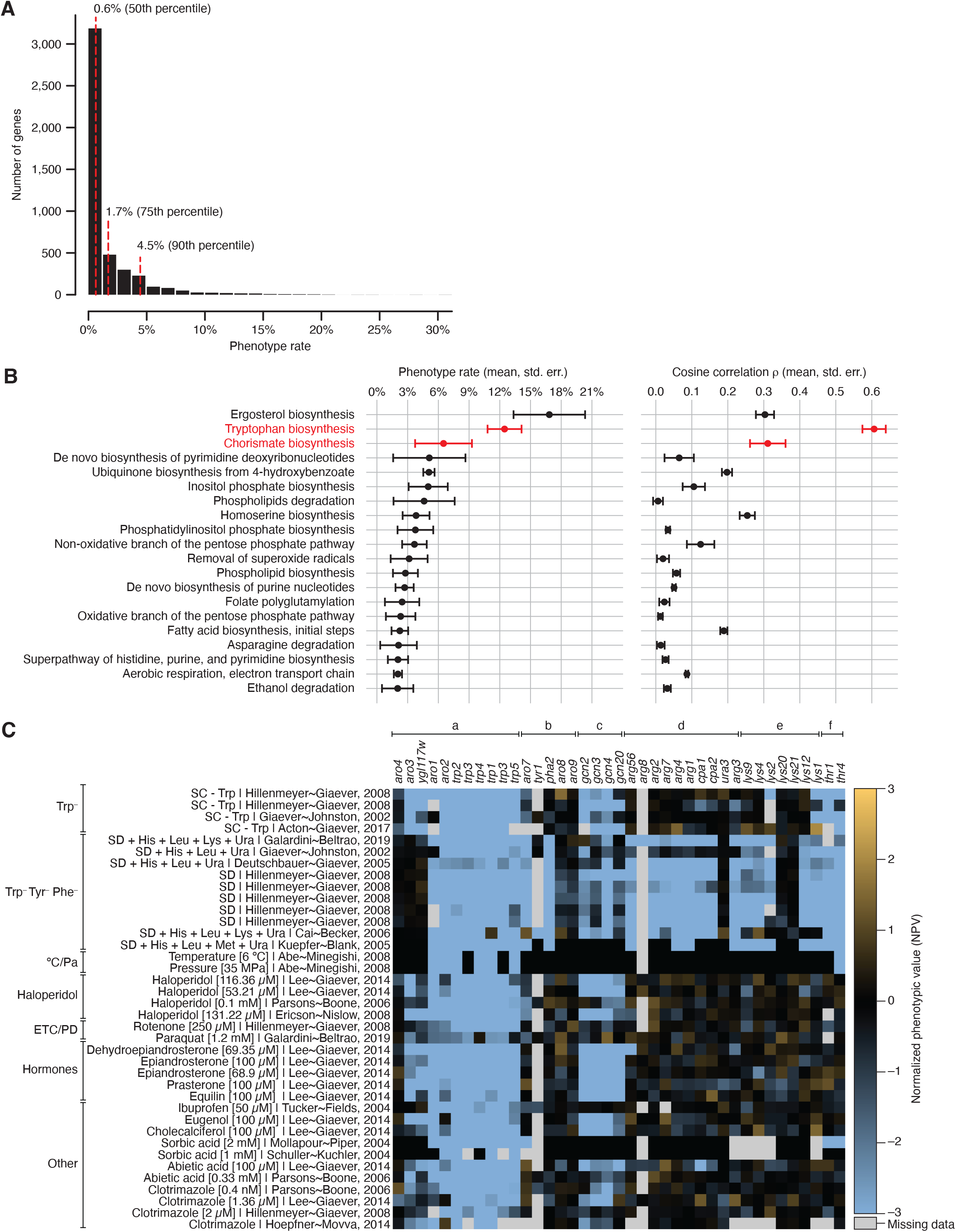
Tryptophan biosynthesis is essential for resistance to a wide range of chemical compounds. (**A**) Distribution of phenotype rates for all genes in Yeast Phenome. (**B**) The biosynthesis of tryptophan and its precursor chorismate are 2 of the top 3 biochemical pathways with the highest phenotype rate. (**C**) Mutants involved in the biosynthesis of tryptophan (*trp1–5*) and chorismate (*aro1–4*), but not other amino acids, share sensitivity to tryptophan-depleted media, low temperature, high pressure and a wide range of chemical compounds. The heatmap shows NPVs for a set of mutants (columns) in a sample of screens (rows). Mutants (columns) are organized by pathway and include: (a) biosynthesis of chorismate and tryptophan; (b) biosynthesis of tyrosine and phenylalanine, (c) the general amino acid control (GAAC) pathway, (d) biosynthesis of arginine, (e) biosynthesis of lysine, (f) biosynthesis of threonine. Screens (rows) are organized by tested condition and include growth in: tryptophan-limited media (Trp^−^), media limited for multiple amino acids, including tryptophan, tyrosine, and phenylalanine (Trp^−^ Tyr^-^ Phe), exposure to low temperature and high pressure (°C/Pa), exposure to haloperidol (Haloperidol), exposure to rotenone and paraquat (ETC/PD), exposure to human hormones (Hormones), exposure to other chemical compounds (Other).

Another potential source of experimental noise in YKO data are secondary mutations (i.e., “suppressors”) that arise spontaneously as adaptations to gene loss and may interfere with the correct assignment of genes to functions. To measure the impact of such strain evolution, we compared different versions of the YKO collection, as well as strains with and without evidence of secondary mutations (note S2; fig. S3). We found that secondary mutations increase the relative risk of incorrect gene-to-function assignment by no more than 3% and, therefore, are unlikely to impede the use and interpretation of YKO data (note S2).

High quality knock-out phenotypes provide strong experimental evidence of gene function and have long been exploited to identify key players in major biological pathways. A multidimensional phenotypic profile, i.e., a vector of binary or quantitative phenotypic values associated with a given gene, is even more powerful at predicting gene function because it enables more robust comparisons of known and unknown genes, and facilitates transfers of knowledge through “guilt-by-association” (*6–10*). We asked how well gene function can be predicted by phenotypic profiles assembled in Yeast Phenome relative to other sources of functional information, such as gene expression, genetic interactions and protein-protein interactions (Materials & Methods). In each dataset, we ranked all gene pairs by their profile similarity and performed a precision-recall analysis using membership in the same functional group (a protein complex, a biochemical pathway, or a moderately specific Gene Ontology biological process term) as ground truth for a functional relationship (Materials & Methods). We found that profile similarity in each dataset is comparably predictive of a functional relationship (area under the precision-recall curve, AUPR = 0.424–0.477; fig. S4A). However, different types of functional relationships are better predicted by different types of biological data (fig. S4B). For example, genes acting in the same biochemical pathway are best predicted by co-expression profiles (AUPR = 0.258), whereas shared membership in the same protein complex is best predicted by similar knock-out phenotypes (AUPR = 0.429). Despite a consistent performance overall, we observed little redundancy between data types such that genes correlated in one dataset were largely uncorrelated in others (fig. S4C). We conclude that each data type provides independent functional information that should be regarded as complementary and analyzed in an integrative manner.

### Yeast Phenome enables novel biological discoveries

Normalized phenotypic values (NPVs), which express a mutant’s phenotype as a standardized deviation from the most typical mutant in the corresponding phenotypic screen, allow us to compare phenotypes across different experiments and identify genes having the greatest impact on cell physiology. We found that virtually all genes have at least one strong phenotype in Yeast Phenome (|NPV| > 3), supporting earlier predictions that no gene is truly dispensable (*8*). Despite this common baseline, the gene-specific phenotype rate, defined as the fraction of screens in which a gene shows a strong phenotype (|NPV| > 3), is highly variable, ranging from ~0% to 31% (mean = 1.8%, median = 0.6%; Fig. 2A). As expected, genes with many phenotypes (top decile, phenotype rate > 4.5%) are more likely to lack a paralogue (odds ratio OR = 2.9, p-value = 6 x 10^-10^), be conserved in higher organisms (OR = 2.7, p-value = 3.8 x 10^-13^) and be annotated to multiple biological processes (OR = 7.4, p-value = 4.6 x 10^-33^) than genes with few phenotypes (bottom decile, phenotype rate < 0.3%). Phenotype rate is also not uniformly distributed across biological processes: genes involved in intracellular trafficking (e.g., intra-Golgi, Golgi-to-endosome and Golgi-to-vacuole transport), pH regulation (e.g., vacuole organization and acidification), lipid metabolism (e.g., ergosterol biosynthesis), transcription, and chromatin remodeling have more phenotypes than expected by random chance (fig. S5A). In contrast, metabolic functions (e.g., transmembrane transport and metabolism of carbohydrates, metal ions and nitrogen compounds) are generally depleted for phenotypes (fig. S5B).

### Tryptophan biosynthesis is essential for resistance to many chemical perturbations

Even though knock-out mutants of most metabolic genes have few phenotypes, we found that biosynthesis of aromatic amino acids is a striking exception and presents one of the highest phenotype rates of all biological processes (fig. S5A). The three aromatic amino acids (tryptophan, tyrosine and phenylalanine) are synthesized from a common precursor, chorismate, via three separate pathways (fig. S6A). However, only genes involved in the biosynthesis of chorismate (*ARO1–4*) and tryptophan (*TRP1–5*) have high phenotype rates (on average, 6.5% and 12.5%, respectively, a 3.8–7.3-fold increase over the mean of all genes; Fig. 2B), whereas tyrosine and phenylalanine biosynthesis genes (*ARO7–9, TYR1, PHA2*) are close to average (1.7%). The phenotype rates of *trp*Δ and *aro*Δ mutants are the second and third highest among 187 biochemical pathways encoded in the yeast genome, following only ergosterol biosynthesis (Fig. 2B). Furthermore, *trp*Δ and *aro*Δ phenotypic profiles are the most highly correlated (cosine ρ = 0.60 ± 0.15 for *trp*Δ mutants, mean ± std. dev.; n = 10 pairs; Fig. 2B), indicating that their phenotypes are likely biologically meaningful and not caused by experimental noise.

As expected, *trp*Δ/*aro*Δ mutants share phenotypes such as the inability to grow on tryptophan-limited media (Fig. 2C, Trp^-^, Trp^-^ Tyr^-^ Phe^-^), at low temperature or under high hydrostatic pressure (Fig. 2C, °C/Pa). Both of these latter conditions are associated with the downregulation of the main tryptophan permease Tat2 and consequent repression of tryptophan uptake (*11*). Interestingly, the vast majority (99%) of *trp*Δ and *aro*Δ phenotypes are sensitivities to 1,138 chemical compounds (NPV < –2), consistent with prior identification of *TRP1–5* and *ARO1–2* as multidrug resistance genes (*8*). The sensitivity of *trpΔ/aro*Δ mutants suggests that these 1,138 compounds modulate tryptophan uptake or metabolism through a direct or indirect mechanism (see Discussion).

Although many compounds causing *trp*Δ/*aro*Δ sensitivity are not easily identifiable because they lack a publicly available name or chemical structure, others are well known chemicals with extensive evidence for a role in tryptophan homeostasis in yeast, rats and other organisms. Examples of such known chemicals are haloperidol, rotenone and paraquat. Haloperidol is an FDA-approved anti-psychotic medication prescribed for the treatment of schizophrenia, Tourette syndrome, bipolar disorder and substance abuse. Long-term haloperidol usage can cause patients to develop tardive dyskinesia (TD), a syndrome of involuntary repetitive body movements such as twitching, shaking and grimacing (*12*). Such movements are greatly reduced by the dietary supplementation of tryptophan in haloperidol-induced rat models of TD (*13*). Rotenone and paraquat are broad-spectrum pesticides that target the electron transfer chain (ETC) and cause oxidative damage. Chronic exposure to both chemicals has been linked to the development of Parkinson’s disease (PD) in mice, rats and humans (*14*). In a manner similar to haloperidol, dietary tryptophan improves the impaired motor functions in rotenone-induced rat models of PD (*15*). The benefits of tryptophan in animals exposed to haloperidol, rotenone and paraquat, along with the sensitivity of yeast *trp*Δ/*aro*Δ mutants to all 3 compounds (Fig. 2C, Haloperidol, ETC/PD), lead us to speculate that these and, potentially, many other *trp*Δ/*aro*Δ chemicals limit the availability of tryptophan in the human nervous system.

Environmental conditions and chemicals causing *trp*Δ/*aro*Δ sensitivity do not impact the growth of other mutants defective in amino acid biosynthesis (e.g., arginine, lysine, threonine; Fig. 2C, d–f). These treatments therefore appear to specifically mimic tryptophan depletion, rather than a general state of amino acid starvation. In wild-type yeast, the availability of all amino acids, including tryptophan, is monitored by the general amino acid control (GAAC) pathway, which senses the accumulation of uncharged tRNAs and upregulates the expression of biosynthetic genes (*16*). Interestingly, GAAC mutants (*gcn2Δ, gcn3Δ, gcn4*Δ and *gcn20Δ*) are sensitive to only ~38% of the conditions that cause *trpΔ/aro*Δ sensitivity (Fig. 2C, c; fig. S6B), suggesting that, under these conditions, the concentration of tryptophanyl-tRNA molecules is indeed decreased and GAAC is required to activate a proper response. In the remaining ~62% of *trp*Δ/*aro*Δ conditions, a functional GAAC is not required for survival, suggesting that tRNA charging is not affected, but that other tryptophan-derived molecules may be limiting (see Discussion).

Overall, the tryptophan biosynthesis pathway appears to be uniquely important for resistance to a wide variety of chemical stresses, some of which may result in decreased tRNA charging. While the specific mechanism for these effects remains unknown, we speculate that *trpΔ/aro*Δ compounds may disrupt the composition and/or fluidity of the plasma membrane, therefore impacting the function of membrane-bound tryptophan permeases (see Discussion).

### Phenotypic profiles organize genes into functional domains

As shown above, phenotypic profiles are powerful tools for identifying functionally similar genes and transfer knowledge through “guilt-by-association” (fig. S4). To gain a global view of gene-gene relationships uncovered by phenotypic similarity, we selected 1,586 genes showing a strong phenotype (|NPV| > 3) in at least 1% of screens and projected the genes on a 2D plane such that their relative distances reflected their phenotypic similarities (Fig. 3A; Materials & Methods). The resulting phenotypic similarity map, annotated with SAFE (*5*), showed that, similar to pair-wise genetic interactions (*17, 18*), knock-out phenotypes organize genes into distinct yet closely connected domains, each enriched for one or more biological processes (Fig. 3A; Materials & Methods).

**Fig. 3:**
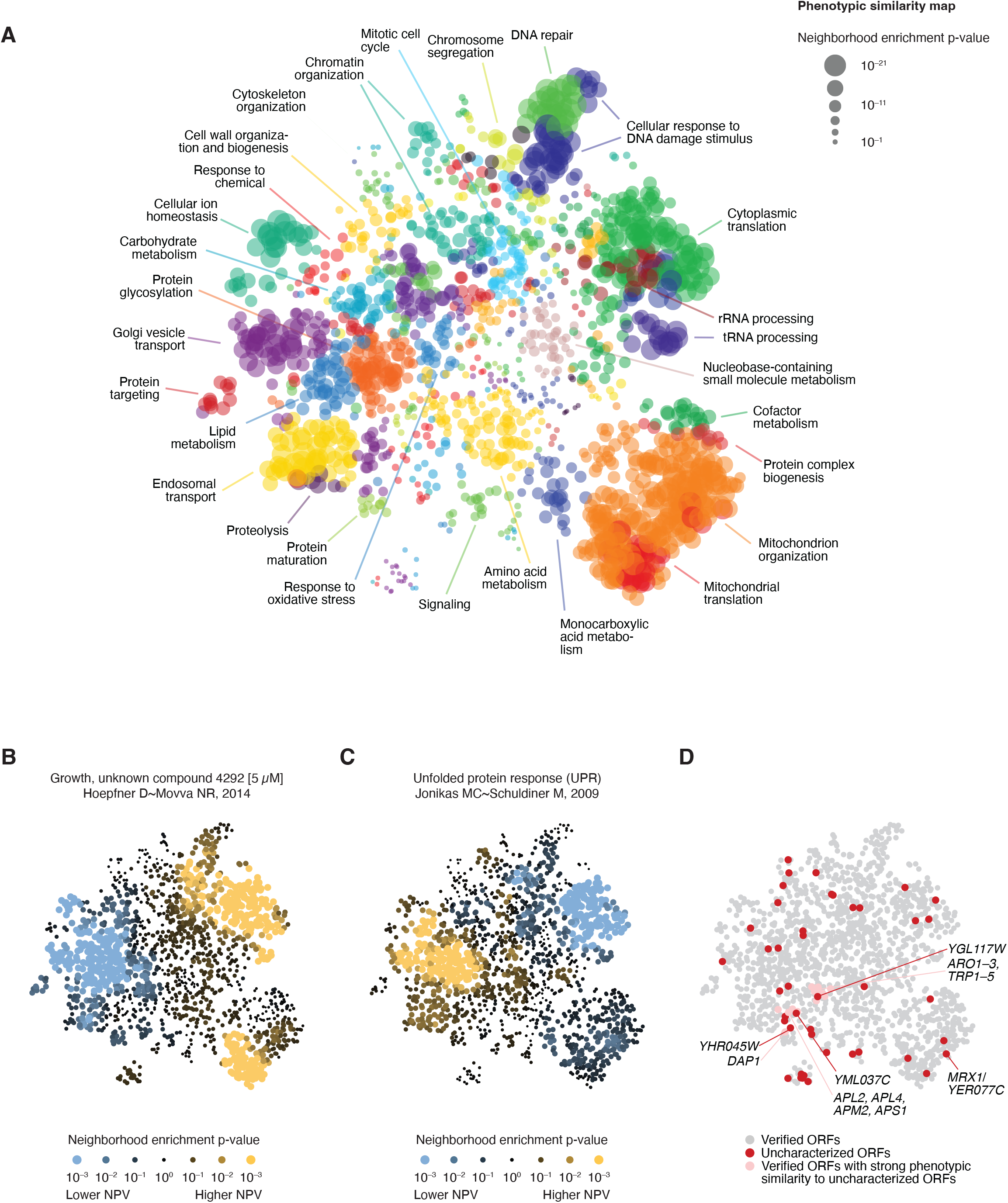
Phenotypic profiles organize genes by function, help interpret novel screens and validate uncharacterized ORFs. (**A**) A phenotypic similarity map was generated by applying UMAP to the phenotypic profiles of 1,586 genes with >1% phenotype rate. The map, where genes with similar phenotypes are placed closer than genes with dissimilar phenotypes, was annotated using SAFE with GO Slim biological process terms. Nodes (genes) are colored based on the GO term with the highest enrichment in their local neighborhoods. The regions with the strongest enrichments are labeled with the corresponding GO terms. (**B–C**) SAFE was used to annotate the map with NPVs from a chemical genomic screen of the unknown chemical compound 4292 (B) and a reporter screen for unfolded protein response (UPR) (C). Nodes (genes) are colored based on the average NPV in their local neighborhood relative to random expectation. (**D**) The phenotypic similarity map shows the distribution of uncharacterized ORFs and suggests hypotheses about their potential functions. Red nodes correspond to 43 uncharacterized ORFs (phenotype rate > 1%, similarity to a verified ORF *ρ* > 0.17). Pink nodes correspond to verified ORFs with strong phenotypic similarity to uncharacterized ORFs.

Importantly, the phenotypic similarity map not only groups genes in a way that reflects their shared function, but also provides a key for interpreting novel or poorly understood phenotypes. For example, the map can be annotated with the chemogenomic profile of an unknown compound to determine which biological processes are required for sensitivity or resistance to the chemical (Fig. 3B). SAFE analysis of one such compound, number 4292 in (*7*), shows that mutants in protein glycosylation, sorting and degradation pathways are sensitive to the chemical, whereas mutants in cytoplasmic and mitochondrial translation are relatively resistant (Fig. 3B). Importantly, the enrichment profile of compound 4292 is a near mirror image of a fluorescent reporter-based screen for unfolded protein response (UPR) (Fig. 3C) that measures the activation of Hac1-regulated genes in response to the accumulation of misfolded proteins in the endoplasmic reticulum (*19*). While the name, molecular target or chemical structure of compound 4292 is not publicly available, the reverse similarity of its enrichment profile to UPR (Pearson R = –0.84, p-value ~ 0) strongly suggests that compound 4292 impairs protein quality control or folding.

### Phenotypic profiles enable annotation of uncharacterized ORFs

The ability of phenotypic profiles to organize genes by function provides an opportunity to validate uncharacterized ORFs and assign novel gene functions. The *Saccharomyces* Genome Database (SGD) estimates that 688 yeast ORFs (~10% of the genome) are currently uncharacterized, meaning that they are likely to produce a protein, as suggested by their conservation in other species, but no such protein product has been experimentally verified in *S. cerevisiae* yet (*20*). Out of the 688 uncharacterized ORFs, 527 ORFs (77%) have at least 10 strong phenotypes in Yeast Phenome (|NPV| > 3) and 46 have robust phenotypic profiles that are predictive of function (phenotype rate > 1%, similarity to a verified ORF *ρ* > 0.17; Fig. 3D, table S3). We found that the top similarities of the uncharacterized ORFs and their positioning on the phenotypic similarity map are highly consistent with preliminary evidence from independent high-throughput experiments, whenever such evidence is available in the literature. For example, *MRX1/YER077C*, which appears to encode a protein localized to mitochondria (*21*) and interacting with the mitochondrial organization of gene expression (MIOREX) complexes (*22*), was most similar to members of the mitochondrial translation machinery and positioned on the map accordingly (Fig. 3D). Another uncharacterized ORF, *YML037C*, mapped next to *APL2, APL4, APM2, APS1* and other members of the AP-1 clathrin-associated adaptor complex (Fig. 3D). This map position is consistent with fluorescence microscopy experiments showing that *YML037C* co-localizes with clathrin-coated vesicles (*21*). To encourage functional annotations of these and other uncharacterized ORFs, as well as verified ORFs without a known function, the Yeast Phenome website provides a set of tools to explore shared phenotypes, verify the mutants’ genomic sequences, and connect to the wealth of information available in other databases (www.yeastphenome.org). As a demonstration of the predictive power of phenotypic similarity, we closely examined two of the uncharacterized ORFs with the highest phenotypic similarity to a verified ORF and tested their predicted functions experimentally.

The first ORF is *YHR045W*, a putative protein of unknown function. Among all mutants in Yeast Phenome, *yhr045w*Δ shows the strongest phenotypic similarity to *dap1*Δ (cosine ρ = 0.59 ± 0.07; Fig. 4A) and localizes next to it on the phenotypic similarity map (Fig. 3D). *DAP1* encodes a heme-binding protein that regulates ergosterol biosynthesis and DNA damage response (*23*). One of the phenotypes shared by *dap1*Δ and *yhr045w*Δ is sensitivity to hydroxyurea, an inhibitor of DNA synthesis: both mutants are among the top 15 hits in ~50% of all genome-wide hydroxyurea screens published to date (Fig. 4A, fig. S8A). We experimentally confirmed the sensitivity of *dap1*Δ and *yhr045w*Δ to hydroxyurea (Fig. 4B). We also examined the *dap1Δ yhr045w*Δ double mutant and found that the two genes are epistatic to one another, showing nearly identical degree of sensitivity to hydroxyurea alone and in combination (Fig. 4B). Furthermore, Dap1 is one of only five known physical interactors of Yhr045w (Fig. 4C).

**Fig. 4:**
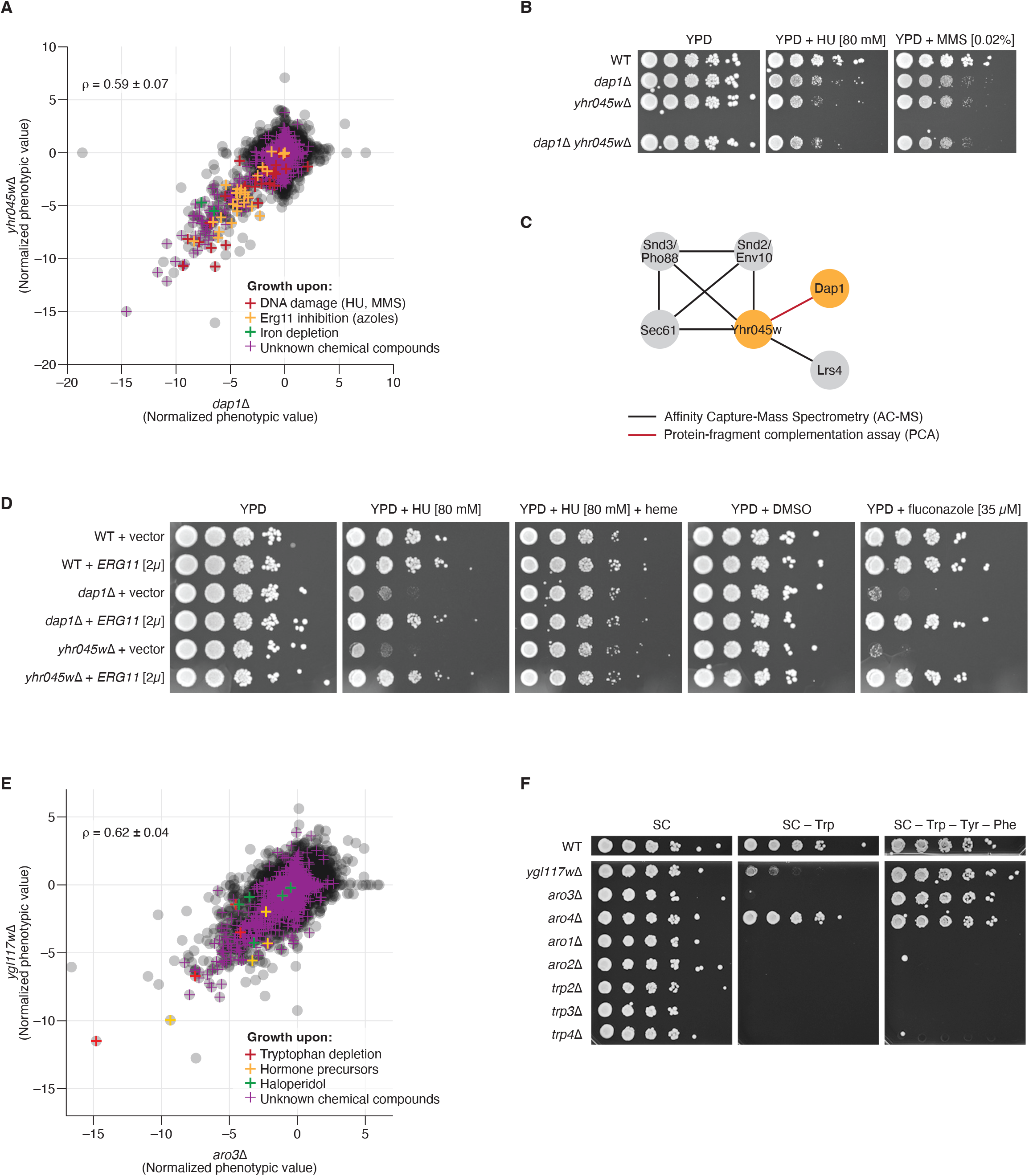
Functional validation of *YHR045W* and *YGL117W*. (**A**) The similarity of the phenotypic profiles of *yhr045w*Δ and *dap1*Δ is shown as a scatter plot of their NPVs. Every grey point corresponds to 1 phenotypic screen. Colored crosses highlight phenotypes suggestive of the genes’ shared function. (**B**) Similar to *dap1*Δ, *yhr045w*Δ is sensitive to DNA damaging agents hydroxyurea and methyl methanesulfonate (MMS). The sensitivity of the *dap1*Δ *yhr045w*Δ double mutant is identical to that of the two single mutants, suggesting that Dap1 and Yhr045w are epistatic to one another. (**C**) Dap1 is one of five known physical interactors of Yhr045w. (**D**) The sensitivity of *dap1*Δ and *yhr045w*Δ to hydroxyurea and fluconazole is suppressed by the overexpression of *ERG11*. The sensitivity of *dap1*Δ and *yhr045w*Δ to hydroxyurea is suppressed by heme supplementation. (**E**) The similarity of the phenotypic profiles of *ygl117w*Δ and *aro3*Δ is shown as a scatter plot of their NPVs. Every grey point corresponds to 1 phenotypic screen. Colored crosses highlight phenotypes suggestive of the genes’ shared function. (**F**) The growth of *ygl117w*Δ is severely impaired in tryptophan-limited conditions (SC–Trp) relative to complete media (SC) but is restored in the absence of all 3 aromatic amino (SC–Trp–Tyr–Phe).

Dap1 is thought to regulate ergosterol biosynthesis by stabilizing Erg11, a member of the cytochrome P450 family that catalyzes the demethylation of lanosterol, an essential intermediate in the ergosterol pathway (*24*). The ability of Dap1 to stabilize Erg11 depends on Dap1’s ability to bind heme, an iron-containing complex that serves as a cofactor in numerous cellular reactions, including Erg11’s demethylation activity (*24*). Consistent with their potential joint role in heme binding and Erg11 stabilization, Yeast Phenome data shows that *dap1*Δ and *yhr045w*Δ are both sensitive to iron depletion and Erg11 inhibition via chemical compounds such as fluconazole and itraconazole (Fig. 4A, D). In addition, large-scale genetic interaction screens have shown that *dap1*Δ and *yhr045w*Δ are both synthetic lethal with a temperature-sensitive *erg11* mutation (*18, 25*), although the overall genetic interaction profiles of *dap1*Δ and *yhr045w*Δ are not significantly correlated (cosine ρ = 0.03 ± 0.11). While the connection between ergosterol biosynthesis and DNA damage is not fully understood, the addition of exogenous heme is able to suppress *dap1*Δ and *yhr045w*Δ sensitivity to DNA damage (Fig. 4D), potentially because excess heme availability bypasses a Dap1-Yhr045w requirement for Erg11 stabilization. Consistent with this hypothesis, overexpression of Erg11 also suppresses *dap1*Δ and *yhr045w*Δ sensitivity to DNA damage and Erg11 inhibitors (Fig. 4D).

To confirm that the phenotypes observed for *yhr045w*Δ are indeed caused by the lack of Yhr045w, we verified its correct genomic sequence in the *Saccharomyces cerevisiae* Genome Variation database (*26*) and complemented its phenotypes with an intact plasmid-borne *YHR045W* (fig. S8B). Taken together, evidence from Yeast Phenome, genetic interaction and protein-protein interaction data, as well as our validation experiments, suggests that Yhr045w acts in cooperation with Dap1 in regulating DNA damage response and ergosterol biosynthesis. To reflect this joint function, we suggest that *YHR045W* be named *DRP1* for “Dap1-related protein 1”.

### Phenotypic profiles enable dissection of complex pathways

The second ORF we chose to characterize is *YGL117W*, a putative protein of unknown function whose phenotypic profile is highly similar to *aro3*Δ (cosine ρ = 0.62 ± 0.04; Fig. 2C; Fig. 3D; Fig. 4E). *ARO3* encodes a 3-deoxy-D-arabino-heptulosonate-7-phosphate (DAHP) synthase, which catalyzes the first step of the chorismate biosynthesis pathway, ultimately producing tryptophan, tyrosine and phenylalanine (fig. S6A). The phenotypic profile of *ygl117w*Δ is as similar to *trpΔ/aro*Δ mutants as they are to one another (Fig. 2C; fig. S7A), strongly suggesting that Ygl117w is a newly identified member or regulator of the pathway. Consistent with this hypothesis, and similar to other amino acid biosynthesis genes, *YGL117W* is upregulated following *GCN4* induction (*27*) and upon amino acid starvation and rapamycin treatment in a *GCN4*-dependent manner (*28*). Furthermore, the promoter of *YGL117W* contains a Gcn4 control response element (GCRE), which is bound by Gcn4 *in vivo* (*29*).

We used the *Saccharomyces cerevisiae* Genome Variation database (*26*) to verify that *ygl117w*Δ mutants in YKO are indeed mutated for *YGL117W*. Furthermore, we experimentally confirmed that, just like *aro3*Δ and all *trp*Δ mutants (but not *aro4*Δ, see below), the growth of *ygl117w*Δ is impaired in tryptophan-limited conditions (Trp^-^; Fig. 4F) and rescued by the expression of a plasmid-borne *YGL117W* (fig. S7B). Interestingly, despite sharing most other phenotypes with the *trp*Δ mutants, *aro3*Δ and *ygl117w*Δ are different from the rest of the pathway in that they can grow when all three aromatic amino acids are missing concurrently (Trp^-^ Tyr^-^ Phe^-^; Fig. 2C; Fig. 4F). Such difference in growth between Trp^-^ and Trp^-^ Tyr^-^ Phe^-^ media is expected for *aro3*Δ, due to Aro3 having functional redundancy with Aro4 (another DAHP synthase) and the feedback inhibition of Aro4 by tyrosine (fig. S6A). However, *aro4*Δ does not mirror this behavior: despite the ability of phenylalanine to inhibit Aro3 activity *in vitro, aro4*Δ exhibits normal growth in Trp^-^ Phe^+^ conditions (Fig. 4F) (*30*). One possibility is that Ygl117w negatively regulates the feedback inhibition of Aro3 by phenylalanine *in vivo* and allows *aro4*Δ to maintain DAHP synthesis in Trp^-^ Phe^+^ conditions (fig. S7C). Overall, to reflect the involvement of Ygl117w in the aromatic amino acid biosynthesis pathway, we propose this gene be named *ARO5*.

### Relationship between phenotypic similarity and intergenic distance

Typically, knock-out phenotypes are attributed exclusively to the deleted gene and interpreted as a reflection of its lost function. However, due to the compact nature of the yeast genome (median intergenic distance = 364 bp, n = 5,864), the deletion of one gene can inadvertently disrupt the accessibility and/or regulation of a neighboring non-overlapping gene. These unintended perturbations, sometimes called neighboring gene effects (NGEs) (*8, 31–36*), are problematic because they can cause changes in expression and/or localization of nearby proteins, and potentially contaminate knock-out experiments with incorrect gene-to-phenotype links. For example, in assigning a new function to *YHR045W*, we verified that *yhr045w*Δ phenotypes were complemented by *YHR045W* but not *YHR042W/NCP1*, a nearby NADP-cytochrome P450 reductase that is also involved in ergosterol biosynthesis and could be indirectly affected by the deletion of *YHR045W* (fig. S8B). While our data indicate that no such perturbation occurs and *yhr045w*Δ phenotypes are indeed due to the loss of *YHR045W*, numerous anecdotal examples of true NGEs have been reported in the literature (note S3).

To systematically measure the extent to which NGEs impact knock-out phenotypes, we used Yeast Phenome data to examine the relationship between phenotypic similarity and intergenic distance for ~782,000 gene pairs located on the same chromosome (Materials & Methods). We found that, consistent with potential NGEs, the phenotypic similarity of immediately adjacent genes was significantly higher than that of all other non-overlapping gene pairs (average cosine ρ = 0.07 vs 0.02, respectively; Kolmogorov-Smirnov test p-value = 1.5 x 10^-248^; fig. S8C). However, to our surprise, excess phenotypic similarity was not limited to adjacent genes: proximal non-adjacent genes, i.e., those located on the same chromosome but not immediately next to one another, also shared significantly more phenotypes than expected (KS test p-value ~ 0.0; fig. S8C). A direct comparison of phenotypic similarity and intergenic distance showed a strong exponential relationship such that, for gene pairs located within ~380 kb of one another, closer proximity corresponded to higher phenotypic similarity, and vice versa (Pearson R = −0.96, p-value = 2.8 x 10^-283^; Fig. 5A). The same trend was observed independently for each chromosome (fig. S9), as well as for multiple unrelated subsets of the Yeast Phenome dataset (fig. S10).

**Fig. 5:**
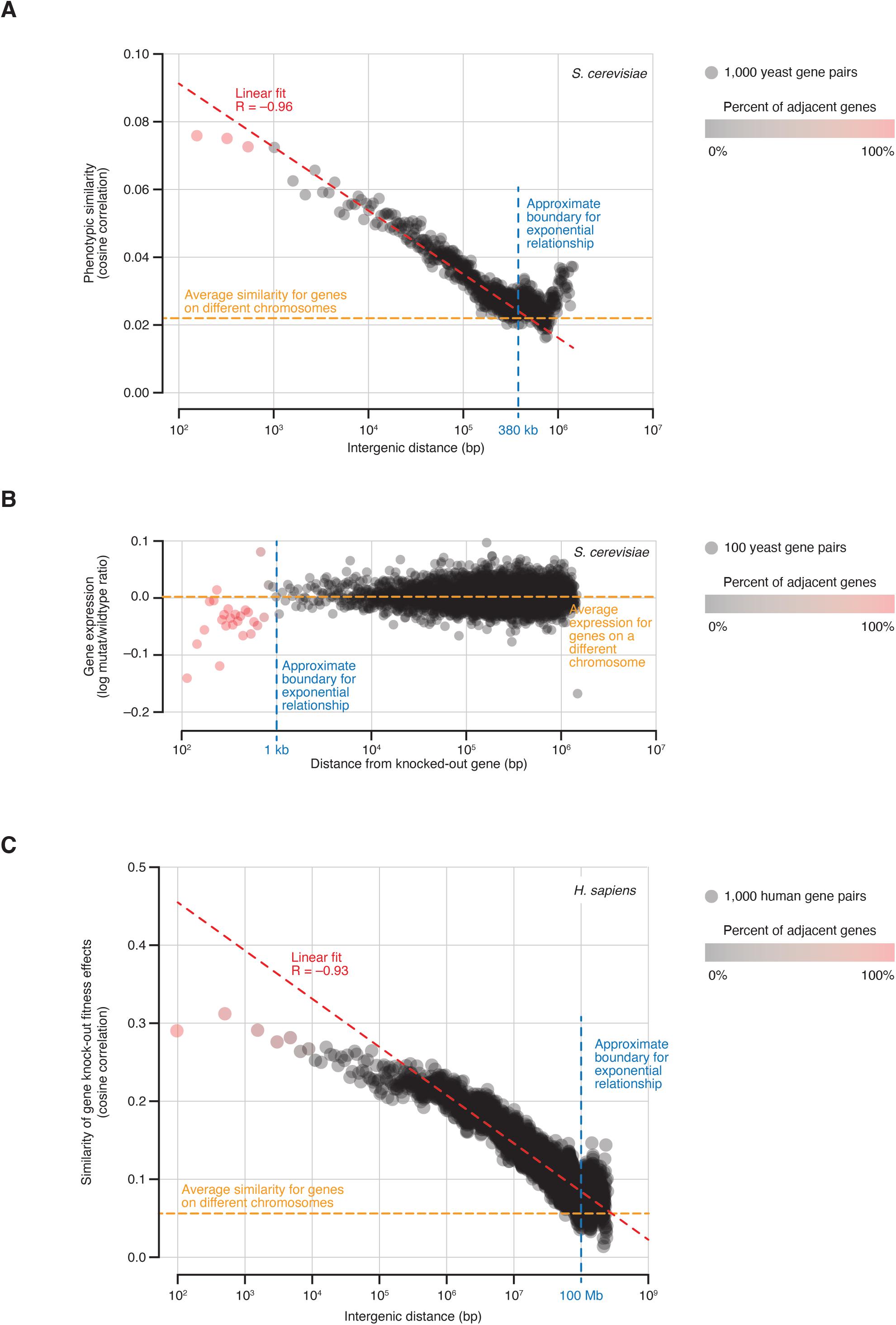
Phenotypic similarity is exponentially related to chromosomal proximity in yeast and human genomes. (A) In the yeast genome, the average similarity of phenotypic profiles decays exponentially as a function of intergenic distance. Gene pairs located on the same chromosome were grouped by intergenic distance. In each group, the average intergenic distance and average phenotypic similarity were computed and plotted on the x and y-axis, respectively. (B) The effect of the knock-out on the expression of nearby genes explains only a small portion of the relationship between intergenic distance and phenotypic similarity. For each knocked-out gene, genes located on the same chromosome were grouped by their distance from the knockout. In each group, the average distance and average change in gene expression in the knock-out strain were computed and plotted on the x and y-axis, respectively. (C) Similar to yeast, the human genome also displays an exponential relationship between intergenic distance and phenotypic similarity. The analysis was done as described in (A). Phenotypic similarity was estimated by comparing gene effects on fitness across ~1,000 cancer cell lines, as measured by genome-wide RNAi and CRISPR loss-of-function screens.

We asked whether the higher phenotypic similarity between proximal genes can be explained by altered gene expression as would be predicted by NGEs (*32–35*). We examined whole-genome transcriptional profiles for ~1,500 knock-out mutants (*4*) and found that genes immediately adjacent to a knock-out are 12 times more likely to change in expression than genes located farther away (0.9% vs 0.08%, respectively; absolute log mutant/wildtype ratio |M| > 1.7, p-value < 0.05; *χ*^2^ p-value = 8.3 x 10^-19^). Most adjacent genes (76%) are downregulated, and, like phenotypic similarity, the magnitude of the effect shows an exponential relationship with chromosomal proximity (Fig. 5B). However, the range of this relationship is much shorter than that observed for phenotypic similarity: on average, only genes located within 1 kb from a knock-out are affected and 92% of these genes are immediately adjacent to the knock-out (Fig. 5B). Such a difference in range between phenotypic similarity and expression effects (380 kb vs 1 kb; Fig. 5A–B) indicates that, while NGEs may be responsible for increased phenotypic similarity among immediately adjacent gene pairs, the phenotypic similarity of proximal non-adjacent genes is likely driven by other factors. One possibility is that this phenotypic similarity reflects a closer functional relationship.

Several studies in yeast and other organisms have reported evidence for chromosomal colocalization of functionally related genes. In yeast, for example, genes that are co-expressed (*37*) or co-regulated by the same transcription factor (*38*), as well as genes encoding members of the same protein complex (*39*) or metabolic pathway (*40*), are more likely to be located nearby on the same chromosome than expected by random chance. To assess the extent to which our observations reflect these known trends, we repeated our analysis after excluding ~186,000 gene pairs with existing evidence of functional co-clustering, as well as paralogous genes arisen from an ancient whole-genome duplication event (Materials & Methods). Interestingly, the exponential decay of phenotypic similarity as a function of intergenic distance was unaffected (fig. S11A), indicating that chromosomal location and biological function have a much stronger connection than previously appreciated.

To confirm that our observations are not due to structural changes in the genome caused by gene deletions, we repeated our analysis in native, unperturbed genomes using co-expression across multiple experimental conditions as a measure of functional similarity (*41*). In agreement with previous reports (*37*), we observed that nearby genes are more co-expressed than genes located farther away or on different chromosomes (fig. S11B). In addition, in a manner consistent with phenotypic similarity, average co-expression decayed exponentially as a function of intergenic distance but affected a much shorter range (up to 10.8 kb, fig. S11B). Similar results were obtained using gene expression measurements obtained via microarray and RNAseq technologies (data not shown).

Finally, we asked whether the relationship between intergenic distance and phenotypic similarity is specific to yeast or is conserved in other organisms. The Cancer Dependency Map Project (DepMap) aims to uncover genetic vulnerabilities in human cancers by systematically inactivating genes in a panel of cancer cell lines and measuring the effect of each gene on cell fitness (*42–44*). Numerous reports have demonstrated that genes sharing similar fitness profiles across cancer cell lines are also likely to share a common function (*45–51*). We examined the similarity of fitness profiles for ~8 million human gene pairs located on the same chromosome and observed the same exponential relationship with intergenic distance as in yeast (R = –0.93, p-value ~ 0.0; Fig. 5C). This relationship, which extends as far as 100 Mb, strongly suggests that, despite differences in genome size, compactness, complexity and perturbation technologies, yeast and human genomes share one fundamental property: genes are not randomly distributed across the genome but positioned relative to one another in a way that reflects their function.

## DISCUSSION

It is commonly assumed that the limiting factor for understanding a biological system is the lack of data or, in some cases, the lack of the right data. Baker’s yeast *Saccharomyces cerevisiae* is a great example of how inaccurate this assumption might be: online repositories and the literature are overflowing with data, yet our understanding of the yeast cell as a complete system is still in its infancy. One reason for such a discrepancy between expectation and reality is that data alone are not sufficient to generate knowledge. To be useful, data must generate hypotheses and, to do so, data must be discoverable, understandable and, most importantly, usable in the context of other types of data (*52*). Yeast Phenome was created to empower integration and re-usability of systematic phenotypic screens of the yeast knock-out collection and fuel the generation of testable hypotheses. By aggregating, annotating and harmonizing all available YKO experiments, we have produced an essential dataset for scientists interested in connecting genotypes to phenotypes, predicting gene function, identifying drug targets, understanding the functional principles of genome organization, testing causal inference methods, and answering many other outstanding questions in the systems biology of yeast and other organisms.

Yeast Phenome incorporates and considerably extends all previous efforts to aggregate yeast knock-out data (*8, 53–57*). In its size, scope and depth of information, Yeast Phenome rivals many human biobanks that aim to facilitate integrative analyses of human biology by linking genomes, phenomes and environomes for hundreds of thousands of individuals worldwide (*1*). However, unlike natural populations, where the effect of a variant on gene function must be predicted from sequence and its contribution to a phenotype must be inferred from statistical associations, a knock-out screen provides a direct measurement of every gene’s causal effect on a phenotype. While in our current work we focused on complete loss-of-function phenotypes, data libraries similar to Yeast Phenome can be created for phenotypes caused by partial loss-of-function, gain-of-function, dosage-modulating and point mutations for which genome-wide collections are already available (*58–65*). As part of our aggregation and annotation efforts, we assembled 7,011 screens of the yeast heterozygous diploid knock-out collection, which capture gene dosage and haploinsufficiency effects on an unprecedented scale. Due to the need to interpret haploinsufficient phenotypes differently from those of loss-of-function mutations, we omitted heterozygous diploid screens from our analyses but are making the dataset available for download and investigation (www.yeastphenome.org; note S4).

We have shown that Yeast Phenome promotes novel hypotheses and a better understanding of cellular biology. One of the advances enabled by Yeast Phenome is the discovery that over 1,000 chemical compounds, including several FDA-approved drugs, limit the intracellular abundance of tryptophan (Fig. 2). While *TRP* and *ARO* genes have been previously linked to multidrug resistance in yeast (*8*), the diversity of Yeast Phenome data provides unprecedented insight into a possible mechanism and its relevance to other organisms. It would be tempting to speculate that the compounds eliciting *trpD/aroD* sensitivity bind and inactivate one or both tryptophan permeases (Tat1 and Tat2), therefore inhibiting tryptophan uptake and making the cell dependent on its biosynthesis. However, the chemical structures of *trpD/aroD* compounds are vastly diverse and their known modes-of-action range from rotenone (a mitochondrial complex I inhibitor) and clotrimazole (an ergosterol biosynthesis inhibitor) to ibuprofen (a non-steroid anti-inflammatory drug) and dehydroepiandrosterone (a human hormone precursor). Such diversity is inconsistent with a direct biochemical interaction with a tryptophan permease or any other protein. A more likely scenario is an indirect effect whereby chemical compounds interfere with tryptophan uptake by changing the structure, composition or fluidity of the plasma membrane. In support of this hypothesis, ibuprofen has been shown to electrostatically adsorb and then hydrophobically insert into phospholipid bilayers in a dose-dependent manner *in vitro* (*66*). Physical perturbations that cause *trpD/aroD* sensitivity (low temperature and high hydrostatic pressure; Fig. 2C) are also known to affect membrane fluidity (*67*). Furthermore, most *trpD/aroD* mutants are synthetic lethal with *erg2–6D* mutants (*18*) which are defective in the production of ergosterol, a primary component of yeast membranes and a regulator of membrane fluidity.

The plasma membrane hosts numerous biomolecules, including sensors, transporters and enzymes, whose function is sensitive to changes in membrane fluidity (*68*). Therefore, it is currently unclear why, relative to all other bioprocesses, tryptophan uptake would be so prominently impacted by membrane perturbations. It is possible that the cell is uniquely sensitive to small changes in tryptophan abundance because tryptophan is the largest, rarest and most energetically expensive of all amino acids (*69*). Furthermore, tryptophan is the only source of *de novo* NAD+ synthesis and may indirectly regulate many metabolic reactions (*69*). In higher organisms, including humans, tryptophan is the precursor of important neuroactive molecules such as serotonin, melatonin, kynurenine, xanthurenic acid and quinolinic acid (*69*), and has been implicated in modulating the ability of tumor cells to evade immune surveillance (*70*). Since human cells are unable to synthesize tryptophan and rely completely on dietary intake, the intracellular availability of tryptophan is determined entirely by the regulation of its transport across membranes. The identification of ~1,000 chemical compounds that may impact such transport will likely be useful in the investigation of neurological diseases and immunooncology.

Another discovery enabled by Yeast Phenome is the exponential relationship between phenotypic similarity and physical proximity among genes located on the same chromosome (Fig. 5). This relationship strongly suggests that genes are not randomly scattered throughout the genome but tend to organize by function. Evidence of co-clustering gene groups has long been available in yeast and other organisms (*37, 71–73*). For example, the major histocompatibility complex (MHC) comprises 20–100 related genes located in the same chromosomal region in most vertebrates (*74*). Our analyses of Yeast Phenome and human DepMap data indicate that this phenomenon is not limited to isolated blocks of functionally similar genes, such as the MHC complex. We show that the relationship between gene position and function is much more continuous and long-ranging than previously appreciated (380 kb and 100 Mb in yeast and human genomes, respectively).

One possible explanation for the pervasive genomic co-localization of functionally related genes is the need to efficiently store and access genetic information within the cell nucleus. Given the complexity of DNA packaging and the energetic costs likely associated with selective access to specific DNA regions, it may not be surprising the genes often accessed together are positioned nearby. Another possible explanation is that physical proximity among functionally related genes has evolutionary advantages for maintaining favorable combinations of alleles. It has been proposed that, when two alleles share a genetic interaction (i.e., their joint effect on fitness is greater than the sum of their individual effects), natural selection should act to preserve the successful haplotype and suppress recombination between the two loci (*73, 75*). Given that functionally related genes are strongly enriched for genetic interactions (*17, 18*), it is possible that their relative genomic positions are under selective pressure to reduce recombination rate and enhance genetic linkage by minimizing physical distance.

A recurrent theme that emerged from our analyses is the importance of examining phenotypic profiles in addition to individual gene-phenotype pairings. A phenotypic profile, intended either as a set of phenotypes associated with a gene or as a set of genes associated with a phenotype, is a powerful tool for investigating a biological system because it is quantitative, comprehensive, and robust to noise. This global perspective is often missed by studies that focus on characterizing only the strongest hits from a loss-of-function screen or, in a largely similar manner, only the most statistically significant variants from a genome-wide association study (GWAS). It is becoming increasingly clear that great value can be derived from examining all genetic variation linked to a trait and all traits linked to a genetic variant, regardless of their significance against an arbitrary threshold.

## ACKNOWLEDGEMENTS

We thank Manuel Hotz, Adam Waite, David Hendrickson, Sean Hackett and Daniel Gottschling for reading the manuscript and providing critical feedback. We also gratefully recognize the generosity of 149 scientists who accepted our invitation and shared unpublished versions of their published screens with Yeast Phenome: Fumiyoshi Abe, Rebecca Abergel, Joaquín Ariño, Prakash Arumugam, Choowong Auesukaree, Simon Avery, Naama Aviram, Alan Bakalinsky, Jeffrey M. Becker, Mohammed Bellaoui, Ajay Bhat, Ron Blackman, Mark R. Bleackley, Javier Botet, Linda Breeden, Grant W. Brown, Anders Bystrom, Steven D. Cappell, Anil Cashikar, Xin Chen, Alessandra Chesi, Leah Cowen, Bobbiejane Culbertson, Valerie Culotta, Bertrand Daignan-Fornier, Scott E. Devine, Henrik Dohlman, Lucília Domingues, Ana Alexandra Mendes Ferreira, Stan Fields, Hernan A. Flores-Rozas, Florian Freimoser, Florian Frohlich, Katsuhide Fujita, Jennifer Gallagher, Imelda Galvan-Márquez, Austen Ganley, David Garfinkel, Susan Gasser, Flaviano Giorgini, Aaron Gitler, Ashkan Golshani, Ramon Gonzalez, Daniel Gonzalez-Ramos, Steven Gorsich, Christopher Grant, Jose M. Guillamon, Sandra Malmgren Hill, Takashi Hirasawa, Dominic Hoepfner, Hans Hombauer, Ambro Van Hoof, Anita Hopper, Dongqing Huang, Sidong Huang, Angelina Huseinovic, Trey Ideker, Yoshiharu Inoue, Philipp A. Jaeger, Torben Heick Jensen, Linghuo Jiang, Xuejiao Jin, Matt Kaeberlein, Patricia Kane, Jerry Kaplan, Brian Kennedy, Ahmad S. Khalil, Hiroshi Kitagaki, Ahmet Koc, Dimitrios Kontoyiannis, Karl Kuchler, Youn-Sig Kwak, Patricia Lage, Emmanuel Lesuisse, Wen-Hsiung Li, Maciej Lis, Beidong Liu, Kirill S. Lobachev, Tiziana Lodi, Monica Montero Lomeli, John Lopes, Kefeng Lu, Walter Luyten, Ross MacGillivray, Andreas Mayer, Mark McCormick, Michael McEachern, Lydie Michaillat, Liz Miller, Eulalia de Nadal, Tomoyuki Nakagawa, Purity Ngina, Francisco Javier Arroyo Nombela, Lorena Norambuena, Thomas Nystroem, Erin O’Shea, Simone Ottonello, Caryn Outten, Barry Panaretou, Amanda G. Paulovich, Youri I. Pavlov, Gemma Perez, Dina Petranovic, Jeff Piotrowski, Markus Ralser, Dindial Ramotar, Peter J. Roach, Michel Roberge, Roberta Ruotolo, Julian Rutherford, Isabel Sa-Correia, Leona Samson, Brian Chia Cheng San, Raul Garcia Sanchez, Martin Schmidt, Udo Schmidt, Brandt L. Schneider, Roger Schneiter, Maya Schuldiner, Shantanu Sengupta, Alexandre Serero, Minyi Shi, Jun Shima, Kenji Shimada, Himanshu Sidha, Sheena Singh-Babak, Gertien Smits, Michael Snyder, Lars Steinmetz, Elena Stepchenkova, Erin Styles, Peter Svensson, Markus Tamas, Saeed Tavazoie, Miguel Teixeira, Martin Tenniswood, Karin Thevissen, Lubomir Tomaska, Chandra Tucker, Bernard Turcotte, Irem Uluisik, Mojca Mattiazzi Usaj, Lieselotte Vermeersch, Kevin Verstrepen, Claudio De Virgilio, Chris Vos, Christopher Vulpe, Tobias Walther, Benedikt Westermann, Wayne Wilson, Fred Winston, Youming Xie, Satoshi Yoshida.

## Funding

This work was funded in part by Calico Life Sciences LLC. The funding source was not involved in preparing the manuscript or in the decision to submit the manuscript for publication.

## Author contributions

Conceptualization: AB

Methodology: CC, BR, VS, JR, RO, FK, MSL, SH, MS, AB

Investigation: GT, RYW, GK, ES, NT, AB

Visualization: GT, VS, AB

Funding acquisition: KJD, DB, AB

Project administration: AB Supervision: KJD, DB, AB Writing – original draft: GT, CC, RYW, ES, AB Writing – review & editing: GT, CC, RYW, ES, VS, JR, RO, NT, KJD, DB, AB

## Competing interests

Authors declare that they have no competing interests.

## Data and materials availability

All Yeast Phenome data are available at www.yeastphenome.org. Code related to the extraction and processing of published data is available at https://github.com/baryshnikova-lab/yp-data.

## Notes

https://www.yeastphenome.org

### Competing Interest Statement

The authors have declared no competing interest.

