## Supplementary Material for "Global analysis of the yeast knock-out phenome"

**This PDF file includes:**

Materials and Methods
Supplementary Text
Figs. S1 to S11
Tables S1 to S4

### Materials & Methods

#### *Aggregation, annotation and normalization of Yeast Phenome data*

*Criteria for inclusion*

In order to be included in Yeast Phenome, a phenotypic screen of the yeast knock-out (YKO) collection must meet all following criteria:

1) Measure a quantitative (continuous) or qualitative (binary, discrete, categorical) phenotype (or the lack thereof) for at least 1,000 knock-out mutants (Note: 1,000 is a cutoff defined empirically by examining the distribution of tested mutants in an early version of Yeast Phenome);
2) Use any current or past version of the haploid Mat-a (BY4741), haploid Mat-α (BY4742) or homozygous diploid (BY4743) deletion collection;
3) Report data on all tested mutants or just the strongest hits (these may include mutants with the largest effect sizes, highest reproducibility or most confident deviations from wildtype);
4) Be associated with a journal article indexed in Pubmed.

Yeast Phenome does not include:

1) Large-scale screens of other mutant collections (e.g., DAmP strains, temperature sensitive mutants or the prototrophic variant of the knock-out library);
2) Studies that report an arbitrary subset of screen results chosen based on biological interest, rather than signal strength or confidence;
3) Genetic interaction screens, i.e., screens where knock-out mutants are examined under a secondary genetic perturbation.

*Strategy for identifying relevant publications*

To identify phenotypic screens that fit these criteria, we developed a comprehensive search strategy and applied it systematically over a 10-year period (2012–2022). As a starting point, we searched the *Saccharomyces* Genome Database (SGD) for gene phenotypes associated with terms such as “systematic mutation set” and “competitive growth”, and compiled a preliminary list of publications that reported phenotypic screens of the YKO collection. By curating these publications and parsing their citations, we discovered many additional publications reporting YKO screens. Furthermore, we examined the publication records of research labs that have released numerous YKO screens and made sure that we captured all of their publications in this domain. We incorporated relevant YKO screens from existing repositories such as the Yeast Functional Genomics Database (YFGdb) (*57*), the Database for High Throughput Screening hits (dHITS) (*54*), ScreenTroll (*55*) and FitSearch (*56*). We received pointers to potentially relevant papers from BioGRID curators and set up automated PubMed queries for keywords such as “yeast knockout collection”, “yeast deletion collection” and “phenotypic screen”.

*Collecting and organizing screen meta-data*

During the curation process, each publication was associated with a list of screens. Each screen was then annotated with extensive meta-data that capture the type of collection (haploid Mat-a, haploid Mat-alpha or homozygous diploid), the phenotype, the experimental environment (including the growth media), the type of released data (quantitative or discrete) and the source from which the data were obtained (see below). The phenotypes and the environments are recorded using a set of controlled vocabularies, i.e. unique terms chosen to avoid duplications and errors. For example, a chemogenomic screen for hydroxyurea was annotated with the phenotype "growth" and the environment "hydroxyurea". Whenever applicable, phenotypes are linked to reporters, i.e. specific readouts through which the phenotype was assessed (e.g., the phenotype "unfolded protein response" has the reporter "UPRE-GFP" since the activation of the response is measured via GFP expression driven by an UPRE promoter). Environments, especially chemical and physical perturbations, are associated with a dose (e.g., environment "hydroxyurea" with dose "100 mM") and alternative names used in the literature (e.g., "hydroxyurea" and "HU"). Whenever available, chemical compounds are also linked with external identifiers from ChEBI (https://www.ebi.ac.uk/chebi/) and PubChem (https://pubchem.ncbi.nlm.nih.gov/) databases.

*Collecting and organizing screen data*

In addition to meta-data describing the features of the experiment, each screen is linked to its corresponding data, i.e. the list of tested mutants and their phenotypic values. The data are obtained from one of three main sources: the main text (e.g., list of hits reported in a table or figure), the supplementary material (e.g., an Excel table or PDF file) or a website associated with the publication (typical for larger datasets). After retrieving the data, we evaluate how complete they are relative to the experiment described in the publication and determine whether additional data might be available. For example, if the publication describes the measurement of a quantitative phenotype but only reports a binary list of hits, we contact the authors and ask if they could share the original quantitative dataset. Similarly, if the list of tested mutants wasn't released as part of the publication, we invite the authors to share that information as well (see below for the explanation why it is important to report the list of tested strains). Whenever we receive unpublished data from the authors, we ask for explicit permission to upload the data onto www.yeastphenome.org. The scientists who agreed to share data are acknowledged as data contributors on the screen's page, as well as in the section describing the project (www.yeastphenome.org/about/data_contributors/). These additional data, which are more complete and, often, more quantitative versions of published data, are explicitly flagged in Yeast Phenome to emphasize their special status.

*Tested mutants*

By definition, all screens in Yeast Phenome used the YKO collection and tested the vast majority of the ~5,000 non-essential gene knock-outs (the biggest exception is Kemmeren~Hostege, 2014 (*4*) that only tested ~1,500 knock-out mutants). However, our preliminary analyses showed that the composition of the YKO collection varied over time and between labs. Furthermore, many screens had to exclude small subsets of the collection for technical or biological reasons (e.g., failure to transform the strain with a plasmid carrying a fluorescent reporter). As a result of such limitations, the set of mutants tested in one screen could differ from that of another screen by as much as 20%. Such variation can prevent an accurate interpretation of the screen results: it may be unclear whether a particular gene is absent from a screen's hit list because it didn't show a strong phenotype (or didn't validate at re-testing), or because it was never tested in the first place. To address this issue, we did our best at recovering a list of tested strains for as many screens as possible. Whenever the list was not released as part of the original publication, we contacted the authors via email (see above). If the authors were unable to provide the list, we estimated the screen's tested space from the tested spaces of all other screens. The estimate was based on a consensus list, i.e. the list of strains that have been tested in at least 50% of all screens that did declare their tested space (Kemmeren~Holstege, 2014 and Huseinovic~Vos, 2017 were excluded from the estimate because their declared tested space was ~1,500 genes, which is considerably lower than other screens). If a screen reported values outside of the consensus tested space, those values were retained.

*Data extraction and normalization*

For all data, we minimize manual handling to maximize record keeping and reproducibility. The only two operations performed manually are 1) downloading the data from the source; 2) whenever required, converting PDF files to a computer-readable format (text or Excel files). All other data manipulations (quality control, clean up, reformatting, upload to database, etc.) are done programmatically. These typically include fixing typos in gene names, translating gene names into ORFs, excluding ORFs that are currently marked as merged or deleted in SGD, converting phenotype data into digital scores as per convention (see below). For each publication, data manipulations are encoded in a Python-based Jupyter notebook, which is version-controlled and stored in a Github repository (https://github.com/baryshnikova-lab/yp-data).

To facilitate both analysis and interpretation of diverse Yeast Phenome data, we implemented several conventions and normalizations. Phenotypes reported on qualitative (e.g., mild, intermediate, severe) or categorical (e.g., round, elongated, irregular) scales were converted into sets of discrete (1, 2, 3) or binary (0, 1) values. Quantitative and truncated quantitative phenotypes (i.e., those where quantitative values are only reported for the set of mutants considered to be hits) were transformed so that the sign and the magnitude of their values were consistent with how the phenotype and its corresponding experimental condition are defined in their respective controlled vocabularies (i.e., higher values correspond to higher expressions of the phenotype, and vice versa). Since different phenotypes followed dramatically different distributions but were consistently unimodal, we used the mode as a reference to normalize each dataset using a modified z-score transformation.

$$NPV_i = \frac{P_i - P_{mode}}{\sqrt{\frac{1}{N}\sum_{i=1}^{N}(P_i - P_{mode})^2}}$$

Where:

$NPV_i$ = normalized phenotypic value for knock-out mutant *i*
$P_i$ = raw phenotypic value for knock-out mutant *i* (as reported by the original publication)
$P_{mode}$ = mode of the kernel density estimate (KDE) distribution of phenotypic values for all knock-out mutants in this screen (KDE was performed using Gaussian kernels, as implemented in the scipy.stats.gaussian_kde, with a scalar bandwidth of 0.25)
$N$ = total number of knock-out mutants tested in this screen

As a result, all phenotypic values reported in Yeast Phenome can be universally interpreted as standardized deviations from the most typical mutant, which, assuming extreme phenotypes are rare, is also likely to represent wild-type.

*Data availability*
All data, including publications, phenotypes, experimental conditions, genes, normalized phenotypic values (NPV) and phenotypic correlations, are available for browsing, searching, displaying and downloading at https://yeastphenome.org/.
The original phenotypic values, along with the Python code used for data extraction and normalization, are available for browsing and download at the yp-data Github repository (https://github.com/baryshnikova-lab/yp-data).
The addition of new data, as well as updates to existing data (e.g., bug fixes, changes in data interpretation and/or normalization), will occur on a case-by-case basis. Changes will be recorded in the commit history of the yp-data Github repository and appear immediately in the database. The bulk download files will be updated quarterly.

***Data sources***

*Yeast Phenome*
Versions: 2022-02-08 and 2022-10-25
URL: https://www.yeastphenome.org/downloads/
Notes:

1. Similarity of phenotypic profiles was measured for each pair of genes using a bootstrap strategy as described below (*Calculating profile similarity*). The similarity metric was cosine correlation. Screens of gene expression from Kemmeren FC~Holstege FH, 2014 (*4*) were excluded from similarity analyses because they provide data for only ~1,500 knock-out mutants.
2. When comparing phenotypic profiles to genetic interaction, protein-protein interaction and gene expression profiles, genes encoding ribosome components were excluded.

*Genetic interactions*
Publication: Costanzo M~Boone C, 2016 (*18*)
Notes:

1. Similarity of genetic interaction profiles was measured for each pair of query strains using a bootstrap strategy as described below ("Calculating profile similarity"). The similarity metric was cosine correlation. Dubious ORFs, essential genes and genes encoding ribosome components were excluded.

*Protein-protein interactions*
Database: BioGRID (*76*)
URL: https://downloads.thebiogrid.org/File/BioGRID/Release-Archive/BIOGRID-4.3.195/BIOGRID-ORGANISM-4.3.195.tab3.zip
Accessed on: 2021-03-31
Notes:

1. Similarity of protein-protein interaction profiles was measured a bootstrap strategy as described below ("Calculating profile similarity"). The similarity metric was Jaccard index. Dubious ORFs, essential proteins, ribosome components and proteins with fewer than 4 interactions were excluded.

*Gene expression*

Database: SPELL (*41*)
URL: http://sgd-archive.yeastgenome.org/expression/microarray/
Accessed on: 2017-09-24
Notes:

1. Similarity of gene expression profiles was measured using a bootstrap strategy as described below ("Calculating profile similarity"). The similarity metric was cosine correlation. Dubious ORFs, essential genes and genes encoding ribosome components were excluded.

*Transcription factor/target gene mapping*
Publication: Balaji S~Aravind L, 2006 (*77*)

*Protein complexes*
Database: EMBL-EBI Complex Portal (*78*)
URL: http://ftp.ebi.ac.uk/pub/databases/intact/complex/current/complextab/559292.tsv
Accessed on: 2022-11-02

*Biochemical pathways*
Database: Yeast Biochemical Pathway Database (YeastCyc) via *Saccharomyces* Genome Database (SGD) (*20*)
URL: http://sgd-archive.yeastgenome.org/curation/literature/biochemical_pathways.tab
Accessed on: 2022-03-02

*Gene Ontology (GO)*
Database: Gene Ontology Consortium (*79*, *80*)
Accessed on: 2017-12-06
Notes:

1. In the precision-recall analysis of phenotypic profiles, as well the analysis of chromosomal co-clustering, GO was restricted to a list of 295 biological process terms that were previously identified by expert biologists as moderately specific (*81*).

*Multifunctional genes*
Database: *Saccharomyces* Genome Database (SGD) (*20*)
URL: http://sgd-archive.yeastgenome.org/curation/literature/go_slim_mapping.tab
Accessed on:
Notes:

1. A gene was defined as "multifunctional" if it was annotated to 4 or more different GO Slim terms.

*Duplicated gene pairs*
Publication: Byrne KP~Wolfe KH, 2005 (*82*)

*Conserved genes*
Database: The Alliance of Genome Resources (*83*)
URL:
Accessed on: 2022-06-28

Notes:

1. A *S. cerevisiae* gene was defined as conserved if it had an ortholog (i.e., best reciprocal match) in *D. rerio*, *M. musculus*, *R. norvegicus*, *C. elegans*, *D. melanogaster* or *H. sapiens*.

*Uncharacterized ORFs*
Database: *Saccharomyces* Genome Database (SGD) (*20*)
URL:
https://yeastmine.yeastgenome.org/yeastmine/bagDetails.do?scope=all&bagName=Uncharacterized_ORFs
Accessed on: 2022-09-03

*Gene coordinates (S. cerevisiae)*
Database: *Saccharomyces* Genome Database (SGD) (*20*)
URL: http://sgd-archive.yeastgenome.org/curation/chromosomal_feature/SGD_features.tab
Accessed on: 2017-04-03

*Human gene knock-out data*
Database: The Cancer Dependency Map Project (DepMap) (*84*)
URL: https://depmap.org/portal/download/all/ [CRISPR_gene_effect.csv]
Version: 22Q2
Accessed on: 2022-10-10
Notes:

1. The similarity of gene effects was calculated using cosine correlation implemented in deepgraph (*85*).

*Gene coordinates (H. sapiens)*
Database: Gene (NCBI)
URL: https://www.ncbi.nlm.nih.gov/gene
Accessed on: 2022-10-11

***Calculating profile similarity***

To compute robust, outlier-insensitive, gene-gene similarities and corresponding confidence estimates in a computationally efficient and parallelizable manner, we adopted the following bootstrap strategy.

1. Given a data matrix (e.g., the Yeast Phenome dataset), where rows correspond to genes and columns correspond to gene features (e.g., phenotypic screens), we created 100 submatrices by selecting 1,500 columns (~10%) using random sampling with replacement.
2. For each of the submatrices, we computed gene-gene correlations as defined by a chosen similarity metric (e.g., cosine, Pearson or Spearman correlations) using a parallelized implementation by deepgraph (*85*).
3. For each gene pair, we combined the 100 sampled correlations and computed the mean and standard deviation.

***Precision-recall analysis of profile similarities***

**Data for the functional groups (GO biological process terms, protein complexes and biochemical pathways), genetic interactions, protein-protein interactions and gene expression were obtained as described above (see**
*Data sources*). Profile similarities (ρ) were computed as described in the Notes for each data type. Gene pairs were sorted (highest to lowest) by the similarity of their profiles. Recall was defined as the number of functionally related gene pairs with $\rho > \alpha$ (for decreasing values of α). Precision was calculated as the fraction of functionally related gene pairs among all gene pairs with $\rho > \alpha$. In the global analysis, a gene pair was considered functionally related if both genes were co-annotated to the same GO biological process term, protein complex or biochemical pathway. In the stratified analysis, a gene pair was considered functionally related if both genes were co-annotated to a functional group from a specific set (e.g., only GO biological process terms).

Area under the precision-recall curve (AUPR) was calculated as the ratio between the area under the true precision-recall curve and the area under an ideal precision-recall curve which would occur if all functionally related gene pairs were ranked higher than all other gene pairs.

***Phenotype rate analysis***

Phenotype rate for a knock-out mutant *i* was defined as $P_i = N_s/N$, where $N$ is the total number of screens where the knock-out *i* was tested and $N_s$ is the number of screens in which knock-out *i* displayed a strong phenotype, i.e. |NPV| > 3. To avoid biases, 6,112 gene expression screens were excluded from this analysis as they have only tested ~1,500 knock-out mutants.

***Constructing the phenotypic similarity map***

The Yeast Phenome data matrix was restricted to 1,586 genes with at least 1% phenotype rate and 8,260 phenotypic screens with more than 1,500 tested mutants (i.e., we excluded the gene expression data from Kemmeren FC~Holstege FH, 2014). All genes were projected onto a 2D space using the Python implementation of Uniform Manifold Approximation and Projection (UMAP) (*86*) with the following parameters: n_neighbors=10, min_dist=0.75, n_components=2, metric='cosine'.

The UMAP projection was annotated using GO Slim biological process terms (see *Data sources*) and a modified version Spatial Analysis of Functional Enrichment (SAFE) (*5*). Briefly, for each gene, we define a local neighborhood as the set of genes located with a Euclidean distance of *d* from it. In this case, *d* was defined as 7% of the map diameter, i.e. the maximum distance between two genes on the map. At this *d* threshold, a typical neighborhood included 41.03 ± 14.28 genes (mean ± std. dev.). Each neighborhood is tested for enrichment for all GO terms using a standard Fisher's exact test. The GO term with the lowest enrichment p-value is assigned to the gene at the center of the neighborhood.

SAFE was also applied for annotating the UMAP projection with the results of quantitative phenotypic screens. Each phenotypic screen is associated with a set of normalized phenotypic values (NPVs). The NPVs for all genes in a neighborhood were summed to produce a neighborhood phenotypic value $\Sigma_{NPV}$. The NPVs were then randomized 1,000 times and a distribution of random cumulative phenotypic values was produced for each neighborhood. An empirical enrichment p-value was calculated by comparing the observed phenotypic values ($\Sigma_{NPV}$) to the random ones. The p-value can be interpreted as the probability of observing a neighborhood phenotypic value as high or higher than $\Sigma_{NPV}$ by random chance (and the opposite for lower values).

#### *Analysis of phenotypic similarity vs intergenic distance*

Chromosomal coordinates for all genes in the yeast genome were obtained as described in *Data sources*. Intergenic distance was calculated as the difference between the left-most coordinate of the upstream gene and the right-most coordinate of the downstream gene. The relationship between intergenic distance and phenotypic similarity was examined by sorting all gene pairs by their distance, splitting the genes into groups of 1,000 pairs each and computing the average distance and phenotypic similarity within each group.

The upper distance boundary for excess phenotypic similarity (i.e., the inflection point in the relationship between phenotypic similarity and intergenic distance) was estimated as follows: 1) compute a Pearson correlation coefficient between the intergenic distance (on a log scale) and phenotypic similarity for all gene pairs located within a distance *d*; 2) repeat the calculation for a range of *d* values; 3) choose the value of *d* that corresponds to a local optimum of Pearson correlation (i.e., the negative peak with a range of similar *d* values). The same approach was used for analyzing gene expression and human gene knock-out data.

To examine the impact of gene pairs with existing evidence of functional co-clustering, we excluded from the analysis all gene pairs that fit any of the following criteria: 1) gene pairs co-annotated to the same protein complex (see *Protein complexes*); 2) gene pairs co-annotated to the same biochemical pathway (see *Biochemical pathways*); 3) gene pairs co-annotated to the same moderately specific GO biological process term (see *Gene Ontology (GO)*); 4) gene pairs in the $75^{th}$ percentile of co-expression values (cosine $\rho > 0.15$) (see *Gene expression*); 5) genes co-regulated by the same transcription factor (see *Transcription factor/target gene mapping*); 5) duplicated gene pairs (see *Duplicated gene pairs*).

#### *Strain construction*

All single deletion mutants used for validation, as well as an isogenic wildtype control, were taken from the Prototrophic Deletion Collection (PDC) (*58*) (table S4).

The double deletion for *dap1*Δ *yhr045w*Δ (ABY004) was constructed by transforming *dap1*Δ (ABY002) with a NatMX cassette PCR-amplified from pRS40_Nat/pFvL099 (*87*). The amplification primers contained 40 bp of the flanking region (upstream and downstream) of *YHR045W*. The double deletion was confirmed by PCR and selection on clonNat and G418.

Strains overexpressing *ERG11* were constructed by transforming wildtype (ABY001), *dap1*Δ (ABY002) and *yhr045w*Δ (ABY003) with either an *ERG11* multicopy 2μ plasmid or empty vector from the MoBY-ORF 2.0 collection (*64*). All transformations were confirmed by leucine selection.

Plasmids used for *yhr045w*Δ complementation assays were constructed using a pRS412-NatNT2 plasmid backbone. This was constructed by inserting a PCR amplified natNT2 fragment from pFA6a-natNT2 with primers incorporating both 5' and 3' *BglII* restriction sites into pRS412 cut with *BglII* to remove the ADE selection site. The *NCP1* and *DAP1* plasmids for complementation were constructed by amplifying the entire ORF by PCR using oligonucleotide primers incorporating 5' *HindIII* and 3' *XbaI* restriction enzyme sites to allow cloning into the pRS412-NatNT2 vector. The *YHR045W* ORF was synthesized from GenScript and inserted into a pUC57-Mini plasmid. The pRS412-NatNT2 and pUC57-Mini-*YHR045W* plasmids were amplified, digested with *NotI* and *EcoRV* restriction sites and ligated before direct transformation into *E. coli*. All plasmids were confirmed by Sanger sequencing before transformation into the deletion strains.

The plasmid used for *ygl117w*Δ complementation was taken from the MoBY-ORF 2.0 collection (*64*). Transformation was confirmed by leucine selection.

***Spot Assays***

Single colonies were grown overnight in 5 mL YPD culture until saturation. They were diluted to an OD of 1.0 in the morning and washed 3x with water. The cells were then resuspended with 1 mL of YPD. A series of 10-fold serial dilutions were prepared in a 96-well round-bottom plate and spotted using a sterilized 48 or 96-pin pronger onto the appropriate growth medium.

***Media***

*YGL117W validation*
Cells were spotted on either SC, SC–Trp, or SC–Trp–Tyr–Phe agar plates. All media contained Yeast Nitrogen Base (YNB) with 5 g/L ammonium sulfate supplemented with 2% glucose. SC and SC–Trp media were supplemented with 2 g/L SC mix (Sunrise Cat #1300) and 1.98 g/L SC–Trp (Sunrise Cat #1305), respectively. For SC–Trp–Tyr–Phe, amino acids were prepared and added back individually according to the concentrations listed in the description of synthetic complete media (Sunrise Cat #1300), but excluding tryptophan, tyrosine, and phenylalanine.

*YHR045W validation*
1.1x YP agar was microwaved for approximately 3-5 minutes until fully boiled. The solution was then cooled to 50°C in a water bath and 20% glucose stock solution was added to a final

concentration of 2%. 40 mL of media was poured into each empty OMNI plate and chemical compounds were added at the concentrations specified in the table below.

| **Drug (from Sigma Aldrich)** | **Concentration** |
|---|---|
| Heme | 20 µM |
| Hydroxyurea (HU) | 80 mM |
| Methyl methanesulfonate (MMS) | 0.020% |
| Fluconazole | 35 µM |
| Itraconazole | 35 µM |

### Supplementary Text

#### *Note S1 – Reproducibility of YKO screens*

By virtue of its size and robust annotations, Yeast Phenome enables an assessment of knock-out screen reproducibility. An example of an experiment repeated multiple times is growth on glycerol, a non-fermentable carbon source that can only be metabolized via respiration. Inability of knock-out mutants to respire, revealed by their failure to proliferate on rich media containing glycerol as the sole carbon source, has been systematically tested 8 times in 6 different laboratories (table S1). By comparing the results of all 8 screens, we found that each screen reported a similar number of slow growing mutants (NPV < –3) that were deemed respiration deficient (140 ± 42 mutants, mean ± std. dev., n = 8). Importantly, most mutants (57–82%) identified in any one screen were reproduced in at least 5 of the 8 independent replicates, while 0–24% of mutants remained unique to single datasets (fig. S2A).

One possible explanation for genes required for respiration in some but not all screens is that differences in experimental design (e.g., glycerol dosage, type of growth assay, mutant ploidy or mating type) create conditional essentialities. If that is the case, we expect genes unique to any single experiment to share common biological functions as often as genes reproduced in multiple experiments. To test this hypothesis, we used proximity on the genetic interaction similarity (GIS) network as an unbiased measure of shared function. The GIS network connects genes if their mutations have similar effects on the fitness of other mutants (*18*). Without any prior knowledge of gene function, the GIS network places genes relative to one another based on the extent of their functional similarity, producing an unsupervised view of cellular organization spanning multiple levels of resolution, from molecular pathways to organelles (*5*, *18*). We performed Spatial Analysis of Functional Enrichment (SAFE) (*5*) to identify regions of the GIS network that are overrepresented for respiration-deficient mutants from each screen (fig. S2B). We found that the enrichment profiles of all screens were nearly identical to one another (cosine correlation between neighborhood enrichment scores ρ = 0.994 ± 0.003, mean ± std. dev., $n_{pairs}$ = 28) and consistent with respiration, oxidative phosphorylation, and other mitochondrial and metabolic functions (fig. S2B). The enrichment was driven primarily by mutants identified in multiple screens, whereas mutants unique to single experiments scattered randomly throughout the network (fig. S2C). This observation does not support the hypothesis that isolated findings share common biological functions and suggests, instead, that they are likely false positives.

Furthermore, it suggests that, whenever replicate screens are not available, SAFE enrichment profiles could inform our level of confidence in single observations.

#### *Note S2 – Analysis of secondary mutations*

Knock-out collections are built on the principle that genetic loci can be systematically altered, one at a time, while keeping the rest of the genome constant. However, the genomes of knock-out mutants are not expected to remain constant over generations. Depending on time and selective pressure, knock-out mutants may acquire secondary mutations that partially compensate for gene loss and alleviate any corresponding fitness defects. Consistent with this expectation, studies have reported heritable phenotypic heterogeneity within isogenic knock-out populations (*88*), mapped secondary site suppressors using systematic genetic crosses (*89*) and identified a wide range of genomic alterations through whole genome sequencing (*7*, *26*). While knowing that adaptation is a general property of living systems that can be minimized but not eliminated completely, we sought to understand to what extent acquired secondary mutations may affect the interpretation of phenotypes derived from the yeast knock-out collection.

We compared phenotypes across two independently constructed versions of the haploid collection (Mat-a and Mat-α), as well as the homozygous diploid collection produced by mating them. Copies of the collections were housed separately across many laboratories and exposed to vastly different experimental conditions, giving each strain an opportunity to evolve independently from its siblings. Despite the opportunity to diverge, estimates of phenotype rate, i.e. the frequency of strong phenotypes ($|NPV| > 3$) displayed by each gene (see also Fig. 2 and related section in the main text), were correlated across collections (e.g., cosine $\rho = 0.66$–$0.72$; fig. S3A), suggesting that secondary mutations are either rare, reoccur frequently in strains lacking the same gene or have relatively little impact on most phenotypes.

To examine the impact on phenotypes more directly, we asked how often secondary mutations mask existing phenotypes or produce new ones, thus lowering the degree to which a phenotypic profile reflects the function of the deleted gene. Using data from previous investigations (*88*, *89*), we compiled a list of 207 knock-out mutants that show ($n = 103$) or do not show ($n = 104$) evidence of secondary mutations (Materials & Methods). We presented random subsets of this list to two independent examiners and asked them to evaluate the phenotypes of each gene with respect to the gene's known biological function (Materials & Methods). The evaluations provided by each examiner, as well as their consensus, showed no statistical association between evidence of secondary mutations and phenotype-function inconsistency (p-value = 0.98, one-tailed Fisher's exact test; fig. S3B). Indeed, in contrast to expectation, the phenotypes of knock-out mutants carrying secondary mutations were more, not less, likely to agree with the functions of the deleted genes (70% among strains with secondary mutations vs 58% in the control group; fig. S3B). Given these data, we estimate with 95% confidence that secondary mutations increase the relative risk of phenotype-function inconsistency by no more than 3% (relative risk RR = 0.711, 95% CI [0.491, 1.030]; table S2).

The relatively low impact of secondary mutations on the functional interpretation of knock-out phenotypes may be explained by close functional proximity (and therefore high phenotypic similarity) between the two affected genes. Indeed, in cases where secondary mutations have been identified, they often occurred in genes that act in the same biological pathway, protein complex or regulatory response as the deleted gene (*88*, *89*). These observations suggest that secondary mutations are likely to modulate, but not obscure, the phenotypes of the original deletion. Consistent with this hypothesis, knock-out mutants with and without secondary site suppressors show highly similar genetic interaction profiles (Jolanda van Leuwen, personal communication). We therefore conclude that secondary mutations, arising spontaneously during routine laboratory manipulations, should not impede the use and interpretation of phenotypic profiles derived from the yeast knock-out collection.

***Note S3 – Examples of true neighboring gene effects (NGEs)***

Analyses of large protein-protein and genetic interaction datasets identified numerous examples of putative neighboring gene effects (*34*, *36*). Below, we are providing 2 additional examples of NGEs with experimental evidence.

*YGL007W and YGL008C/PMA1 (Porat et al., 2005)*

*YGL007W* is an uncharacterized ORF located ~600 bp upstream of *PMA1* and overlapping its promoter region. *PMA1* encodes an essential H+-ATPase proton pump and is one of the key regulators of membrane potential which enables the transport of amino acids and other molecules, including polyamines (putrescine, spermidine and spermine). The expression of *PMA1* is regulated by the DNA-binding protein Rap1 which recognizes two upstream activating sequences (UASs) located within the coding sequence of *YGL007W* (*90*). As a result, deletion of *YGL007W* causes a ~50% reduction in Pma1 levels and confers increased resistance to spermine (*91*). Expression of a plasmid-borne *YGL007W* does not restore sensitivity to spermine, demonstrating that the *ygl007w*Δ phenotypes are the result of *PMA1* downregulation and not loss of Ygl007w activity (*91*).

*YJL027C and YJL026W/RNR2 (this study)*

Hydroxyurea is a chemical compound that blocks DNA synthesis by inhibiting the ribonucleotide reductase (RNR) complex and depleting the intracellular pool of deoxyribonucleotides (*92*). In Yeast Phenome, the effects of hydroxyurea on growth of knock-out mutants have been assessed quantitatively 13 times by 6 different laboratories with doses spanning 2 orders of magnitude (8.3–200 mM). We compared the relative ranking of mutants across experiments and observed that, as expected, the strongest effects were consistently associated with mutations in DNA replication and repair genes, such as *rad54*Δ, *rad5*Δ and *pol32*Δ, as well as other known modulators of cellular response to replication stress, such as *sod1*Δ, *dhh1*Δ, *lsm1*Δ and *dun1*Δ (fig. S8A). Interestingly, in 5 of the 13 datasets, the most critical factor for survival in hydroxyurea was *YJL027C*, a gene encoding a putative protein of unknown function located 155 bp upstream of *RNR2* and likely overlapping its promoter region. *RNR2* encodes an essential subunit of the RNR complex which is both targeted by hydroxyurea and upregulated in response to DNA damage (*93*). Because of its essentiality, *rnr2*Δ is absent from Yeast Phenome data. However, inability to regulate the expression of *RNR2*, as a result of *YJL027C* deletion, combined with the inhibition of RNR complex activity by hydroxyurea, is likely responsible for the *yjl027c*Δ extreme sensitivity to the compound.

#### *Note S4 – Phenotypic screens of the heterozygous diploid YKO*

The data provided at www.yeastphenome.org and used for analysis in this study include only phenotypic screens of the two haploid YKO collections (Mat-a and Mat-α) and the homozygous diploid YKO collection. However, we also assembled and annotated data derived from phenotypic screens of the heterozygous diploid collection. These data were processed and transformed in the same manner as the haploid/homozygous phenotypic screens (Materials and Methods). The unprocessed input data and processing code for each publication reporting heterozygous screens are provided in the yp-data Github repository (https://github.com/yeastphenome/yp-data). In addition, we're providing the following 3 files in the "Bundle downloads" section of the Yeastphenome.org website (https://www.yeastphenome.org/downloads/bundles/):

1. yp_datasets_het_20221018.tar.gz (tab-delimited) -- A list of heterozygous diploid screens with relevant metadata (explained in the README.txt file).
2. yp_matrix_het_20221018.tar.gz (tab-delimited) -- A gene x screen matrix for cleaned but unnormalized phenotypic values.
3. yp_matrix_het_z_20221018.tar.gz (tab-delimited) – A gene x screen matrix of normalized phenotypic values.

### Figure S1

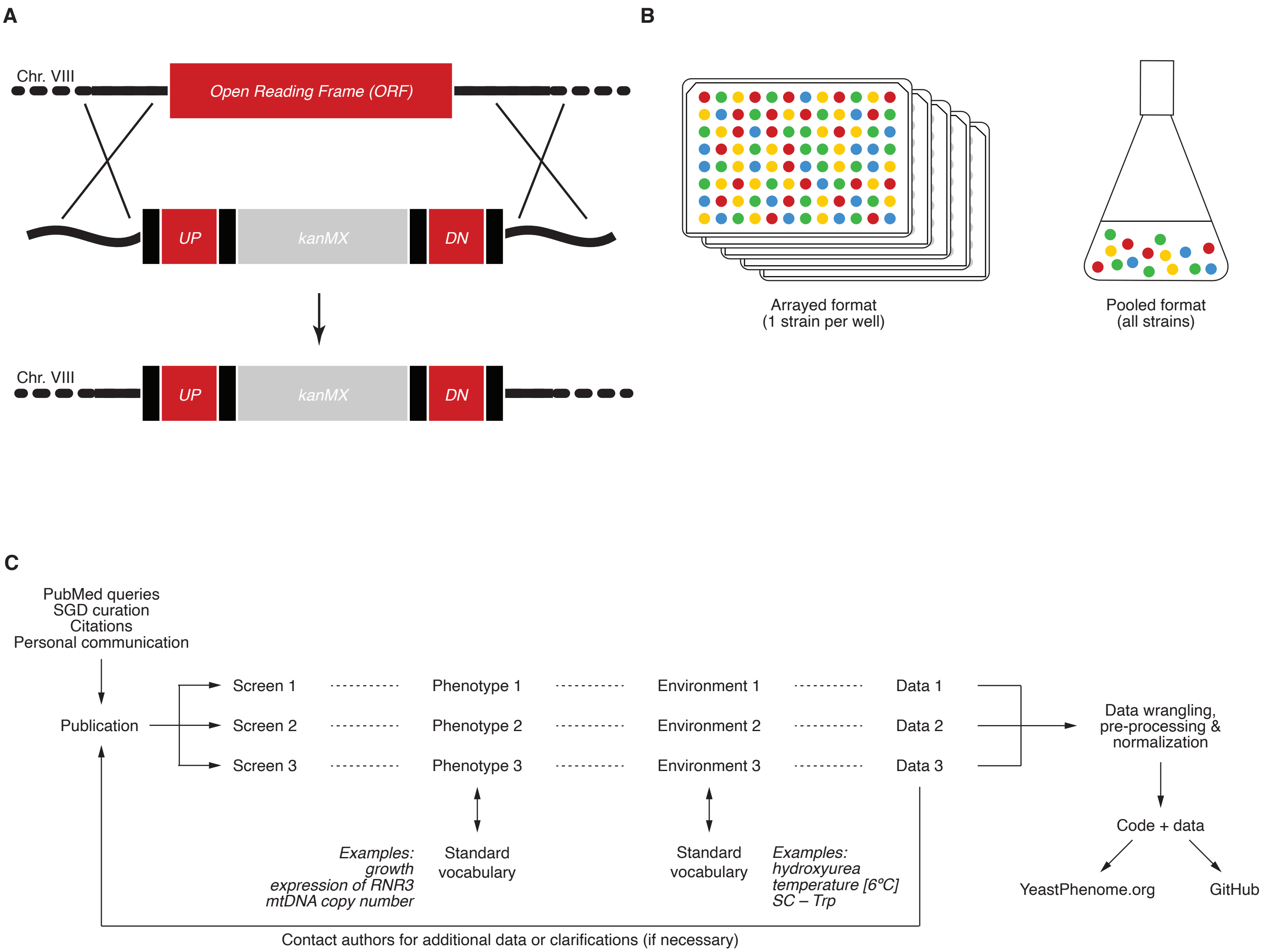


All published screens of the yeast knock-out (YKO) collection were identified, curated, assembled and normalized to enable analysis and integration. (**A**) In the YKO collection, each open reading frame is deleted and replaced via homologous recombination with a selectable marker (kanMX) flanked by locus-specific molecular barcodes (UP and DN), as well as universal sequences that can be used for amplification (black vertical bars). (**B**) Phenotypic screens involving the YKO collection are typically performed in an arrayed or a pooled format. In the arrayed format, each strain is examined independently from other strains by virtue of being grown in a separate well in a 96-well plate and/or as a separate colony on solid media. In the pooled format, all strains are co-cultured together in the same vessel and identified by barcode sequencing or microarray hybridization. (**C**) Publications that report phenotypic screens of the YKO collection were discovered in many ways including PubMed queries, other databases (e.g., SGD), citations and personal communication from yeast researchers. Each publication was associated with a list of screens and each screen was annotated with a set of standard vocabularies, i.e. lists of standardized terms that describe the measured phenotype (e.g., growth, expression of *RNR3*, mtDNA copy number) and the environment or experimental condition in which the phenotype was measured (e.g., growth medium, exposure to a chemical compound, temperature). Each screen was also associated with the corresponding data, which comprise the list of tested knock-out mutants (whenever available) and the list of phenotypic values for each tested mutant (whenever available). These data were cleaned, harmonized and normalized, as described in Materials & Methods. The original and normalized data, as well as the Python code used for processing, were also stored in a database and a GitHub repository (https://github.com/yeastphenome/yp-data).

**Figure S2**

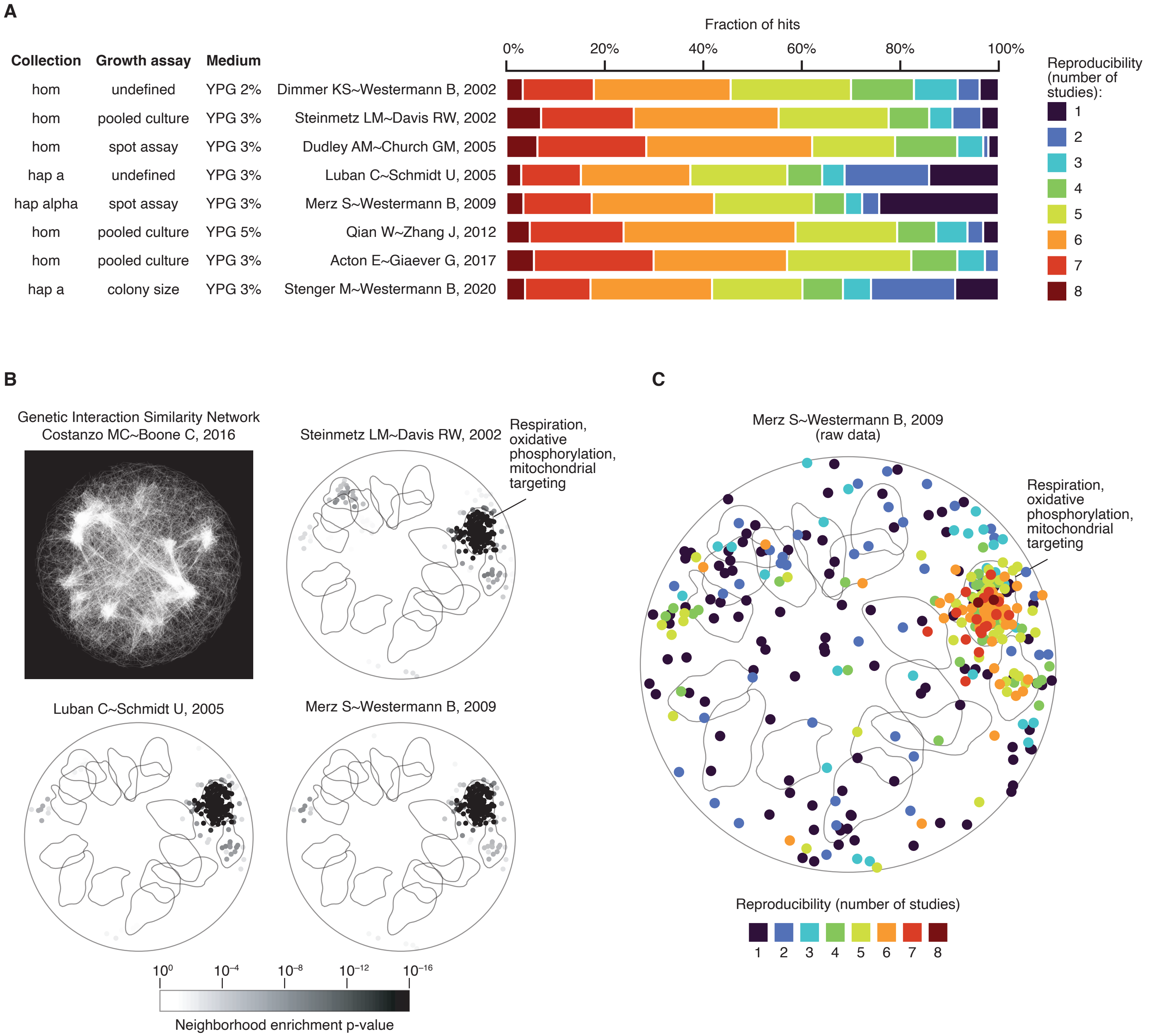


Thanks to its size and meta-data annotations, Yeast Phenome allows to identify similar screens and assess their reproducibility (note S1). (**A**) We identified 8 independent screens of respiratory metabolism (i.e., growth on rich media with glycerol as sole carbon source) using criteria described in Table S1. In each screen, “hits” were defined as knock-out mutants with a strong growth defect (NPV < –3) relative to the most typical mutant in that screen (i.e., mode of all phenotypic values). The fractions of hits identified in one screen and reproduced in 0–7 other screens are shown as stacked bars and color-coded. For example, ~4% of hits reported by the first screen (Dimmer KS~Westermann B, 2002) were unique to that screen (black). In contrast, ~18% of hits were reproduced by 6 or 7 other studies (dark red + red). (**B**) The similarity of the 8 screens for respiratory metabolism was nearly complete when, instead of a gene-by-gene overlap, we compared their SAFE profiles. A screen’s SAFE profile illustrates the statistical association between the identified hits and one or more domains of the genetic interaction similarity network. Visual and quantitative comparisons of the SAFE profiles of the 8 screens (3 of which are shown here) demonstrate that, on a functional level, the sets of identified hits are highly similar to one another and consistently associated with respiration, oxidative phosphorylation and mitochondrial targeting functions. (**C**) Reproducible hits are more likely to associate with relevant biological functions than non-reproducible hits. Nodes of the genetic interaction similarity network (B) that correspond to hits from Dimmer KS~Westermann B, 2002 are represented as dots. The color of each dot indicates the total number of screens in which that gene was identified as a hit. Dark red, red, orange and light green colors indicate hits reproduced in at least 5 of the 8 studies. As expected, these hits are concentrated in the network domain associated with respiration, oxidative phosphorylation and mitochondrial targeting. In contrast, black and dark blue dots indicate genes identified in only 1 or 2 screens. These hits are more randomly distributed throughout the network.

**Figure S3**

**A**

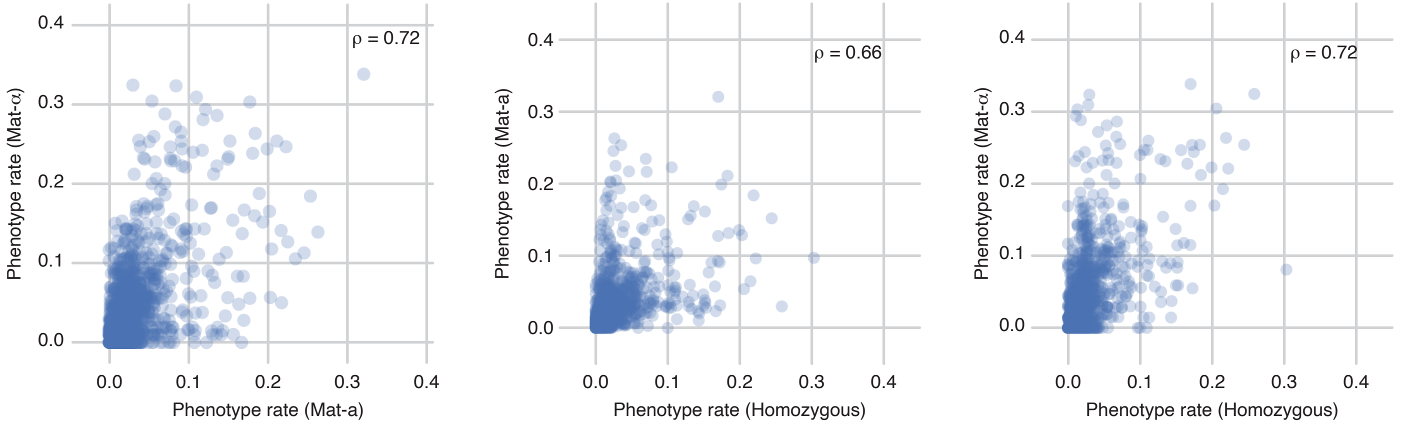


**B**

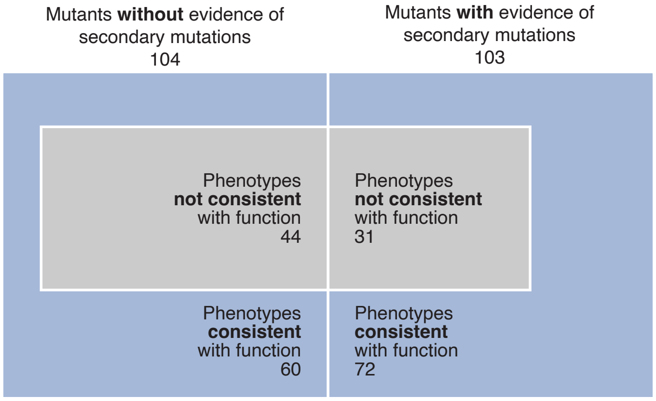


Secondary mutations are unlikely to impact gene-phenotype associations from YKO screens. (**A**) The phenotype rates of Mat-a, Mat-α and homozygous diploid strains mutated for the same genes are generally correlated (cosine correlation ρ = 0.66–0.72), suggesting that secondary mutations are either rare, reoccur frequently in strains lacking the same gene or have relatively little impact on most phenotypes. (**B**) The phenotypic profiles of knock-out mutants with evidence of secondary mutations are more, not less, likely to be consistent with known functions of the knocked-out genes than the phenotypic profiles of mutants without evidence of secondary mutations. Each blue box represents a set of knock-out mutants with (right) and without (left) evidence of secondary mutations (note S2). The grey areas in each box are proportional to the fraction of mutants presenting phenotypes that are inconsistent with the gene's known function.

**Figure S4**

**A**

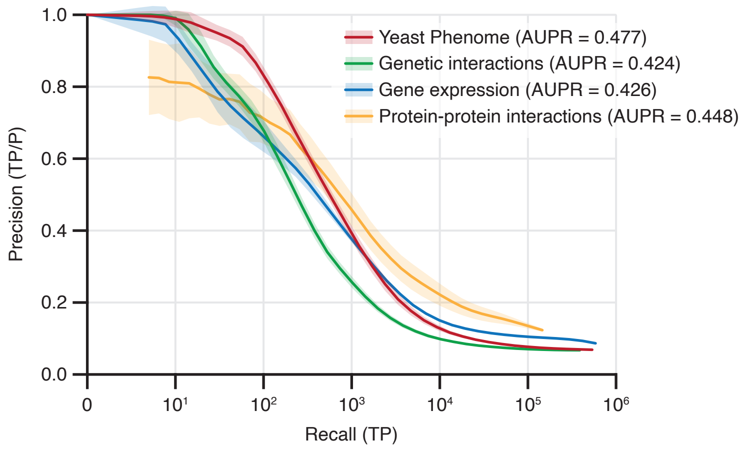


**B**

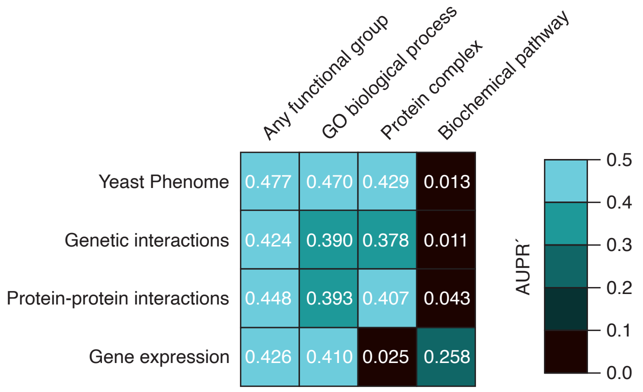


**C**

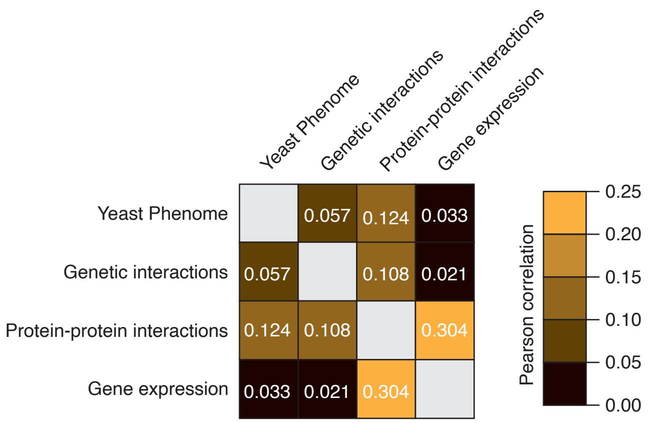


Phenotypic profiles predict functional relationships as accurately as other genome-scale datasets. (**A**) We examined gene-gene similarities using 4 independent data sources: Yeast Phenome, genetic interactions, protein-protein interactions and gene expression (Materials & Methods). For each dataset, we calculated profile similarities using the bootstrap strategy (20 samples, 1,500 features per sample) described in Materials & Methods. In each sample, we ranked gene pairs by their profile similarity (ρ) and computed recall (number of functionally related pairs with $\rho > \alpha$ for decreasing values for α) and precision (the fraction of functionally related pairs among all gene pairs with $\rho > \alpha$ for decreasing values of α). A gene pair was considered functionally related if both genes are co-annotated to the same GO biological process term, protein complex or biochemical pathway. The plot shows the relationship between recall and precision for each dataset. Lines and shaded areas represent the average and standard deviation of precision-recall curves for the 20 samples. Data sources and details about calculating phenotypic similarities, precision-recall and areas under the precision-recall curve (AUPR) are described in Materials & Methods. (**B**) Different types of functional relationships are better predicted by different data types. The heatmap shows areas under the precision-recall curves (AUPRs) computed following the precision-recall analysis described in (A) but using narrower definitions of functional relationship (e.g., only co-annotation to the same protein complex or only co-annotation to the same biochemical pathway). (**C**) Despite an overall consistent performance in functional prediction, we observed little redundancy between data types such that genes correlated in one dataset were generally uncorrelated in others. The heatmap shows Pearson correlation coefficients between gene-gene profile similarity values computed from the 4 different data sources.

**Figure S5**

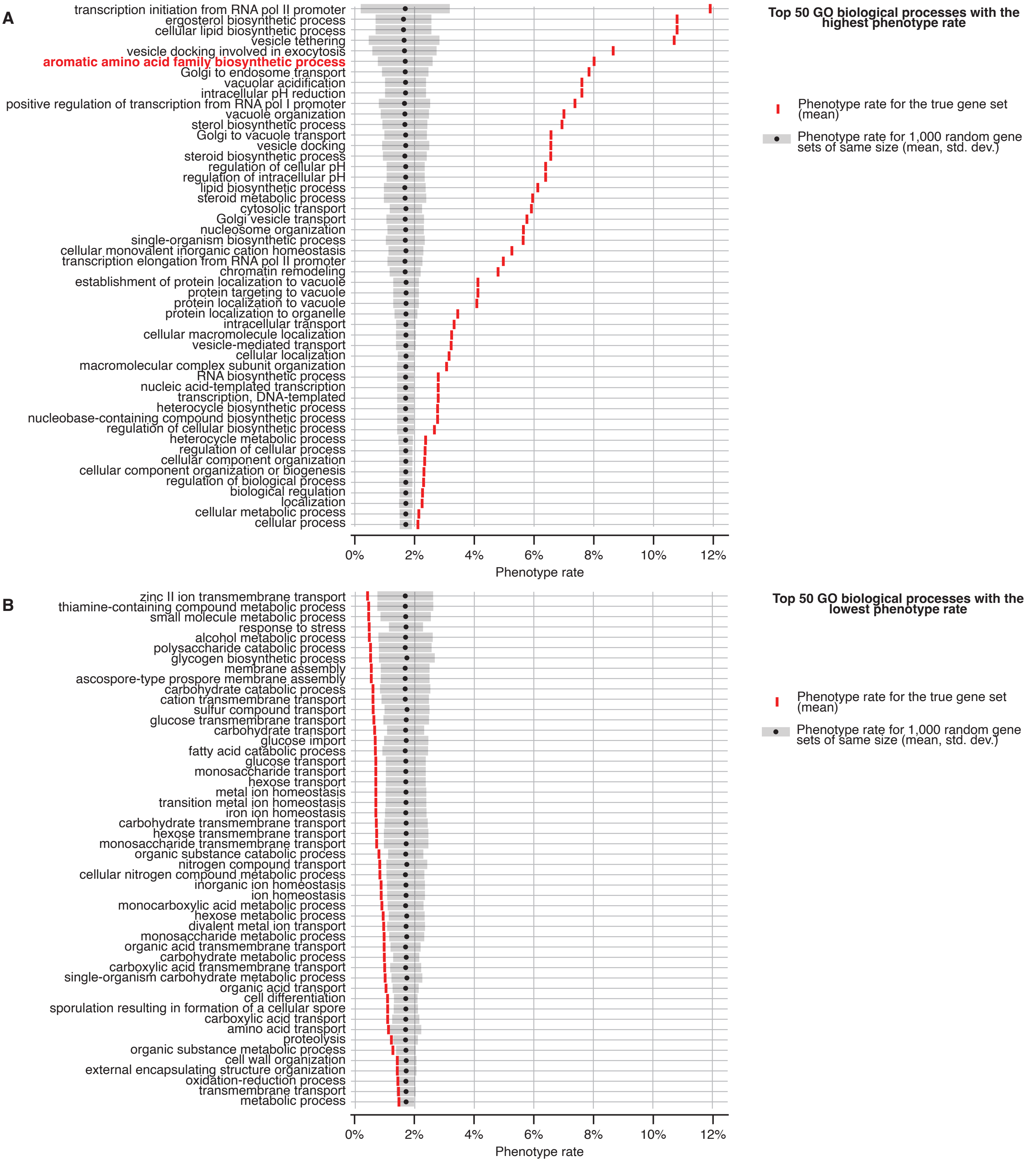


Phenotype rate, defined as the fraction of screens in which a gene shows a strong phenotype (INPVI > 3), is not uniformly distributed across biological processes. An average phenotype rate was computed for each GO biological process with more than 10 genes represented in Yeast Phenome. The true phenotype rate ($P_{true}$, red line) was compared to the average and standard deviation of 1,000 randomly sampled gene sets of the same size ($P_{random}$, black dot and σrandom, grey box, respectively). A z-score for each biological process was computed as $(P_{true} – P_{random}) / \sigma_{random}$. The top 50 biological processes with the highest (**A**) and the lowest (**B**) z-scores are shown. The aromatic amino acid family biosynthetic process (highlighted in red) is the only metabolic process with an elevated phenotype rate.

**Figure S6**

**A**

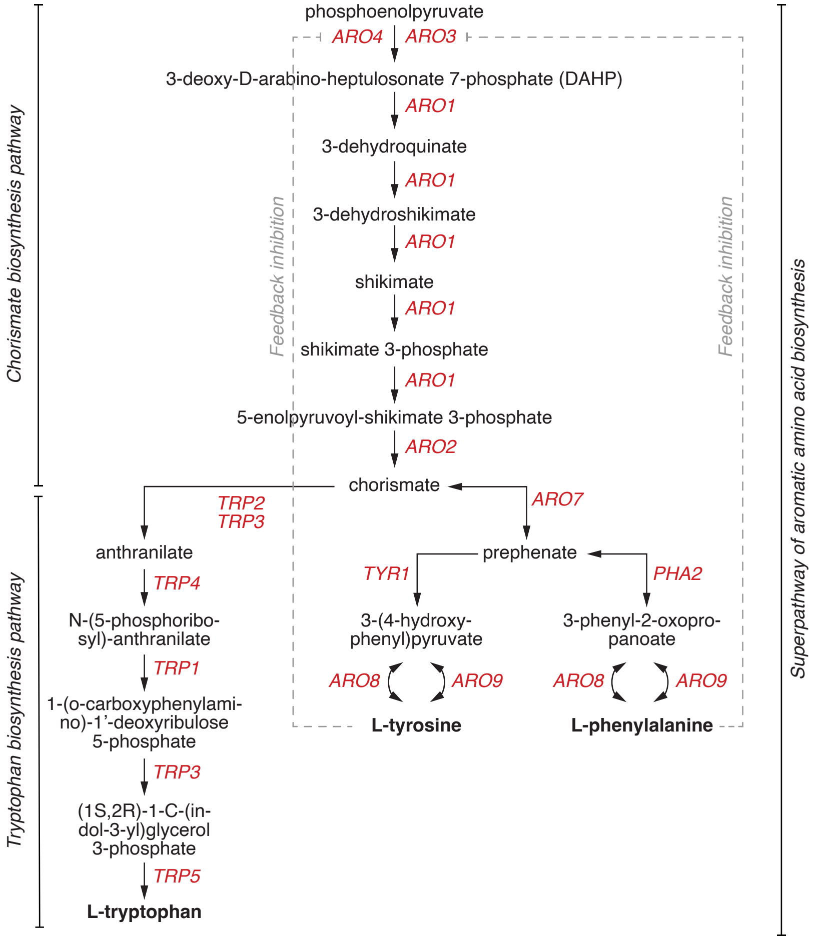


**B**

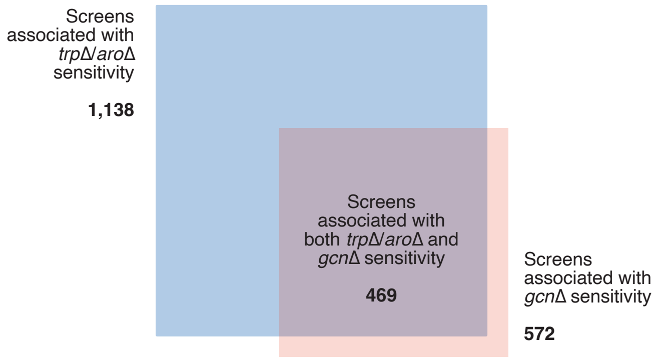


(**A**) The superpathway of aromatic amino acid biosynthesis includes the biosynthesis of tryptophan, tyrosine, and phenylalanine, as well as their common precursor chorismate. Genes encoding enzymes required for each reaction are indicated in red. The feedback inhibition of *ARO4* by tyrosine and *ARO3* by phenylalanine are indicated by the grey dashed lines. (**B**) Mutants impaired in the general amino acid control (GAAC) pathway (*gcn2*Δ, *gcn3*Δ, *gcn4*Δ and *gcn20*Δ) are sensitive to only ~38% of the conditions that cause *trp*Δ/*aro*Δ sensitivity. The Venn diagram shows the overlap between two sets of screens: 1) blue: 1,138 screens where at least 4 of the 8 *trp*Δ/*aro*Δ mutants show impaired growth (NPV < –2); 2) red: 572 screens where at least 2 of the 4 *gcn*Δ mutants show impaired growth (NPV < –2).

**Figure S7**

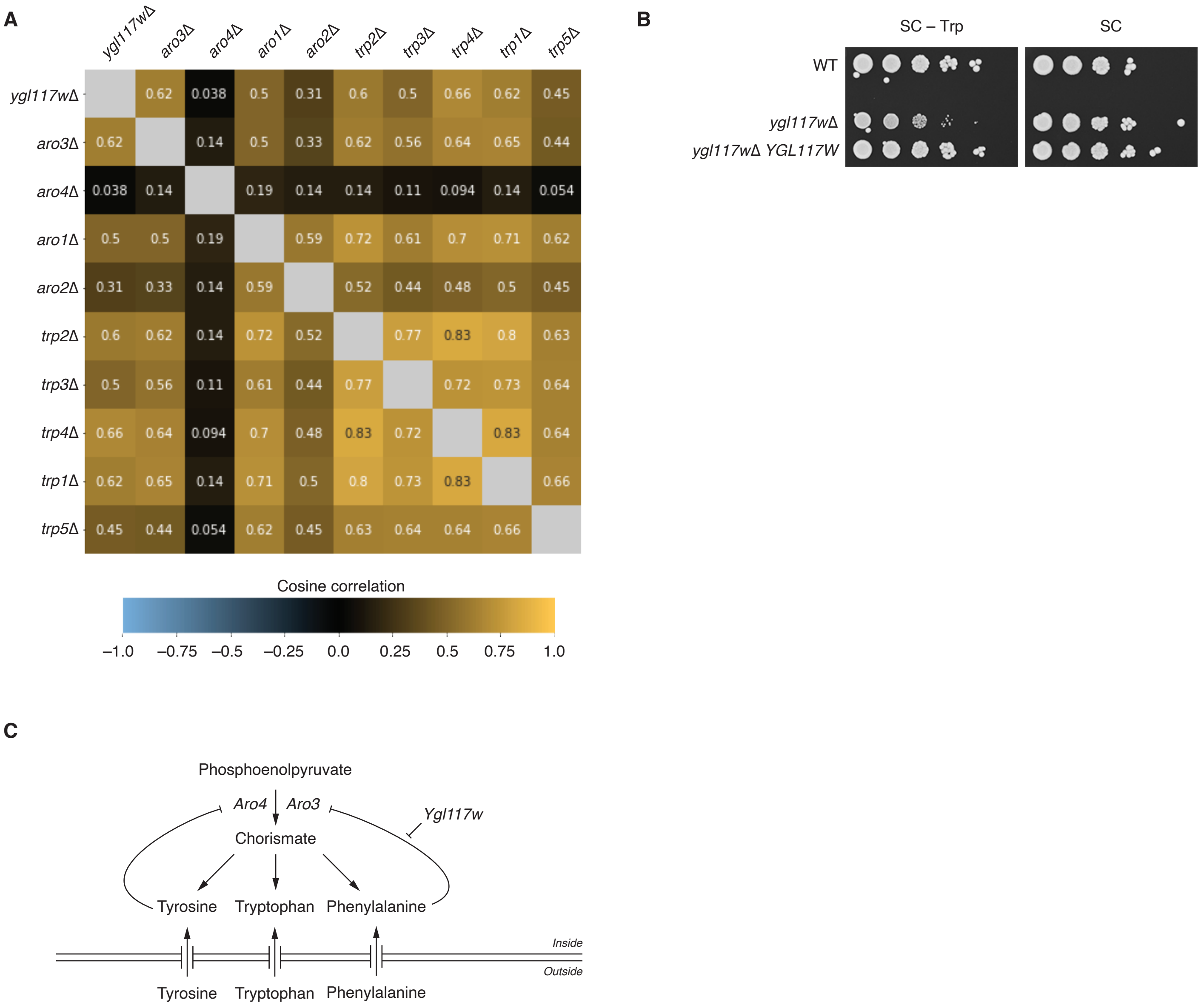


Evidence from Yeast Phenome and validation experiments suggests that *YGL117W* is a novel member or regulator of the aromatic amino acid biosynthesis pathway. (**A**) The phenotypic profile of *ygl117w*Δ is as similar to *trp*Δ/*aro*Δ mutants as they are to one another. Cosine correlations between all pairs of genes were computed using the bootstrap method described in Materials & Methods. (**B**) The growth defect of *ygl117w*Δ on media lacking tryptophan (SC–Trp) is rescued by the expression of a plasmid-borne *YGL117W* (Materials & Methods). (**C**) A model describing the potential role of Ygl117w in the aromatic amino acid biosynthesis pathway. Data in the literature, Yeast Phenome data and our own validation experiments are consistent with the hypothesis that Ygl117w negatively regulates the ability of phenylalanine to feedback-inhibit the DAHP activity of Aro3.

**Figure S8**

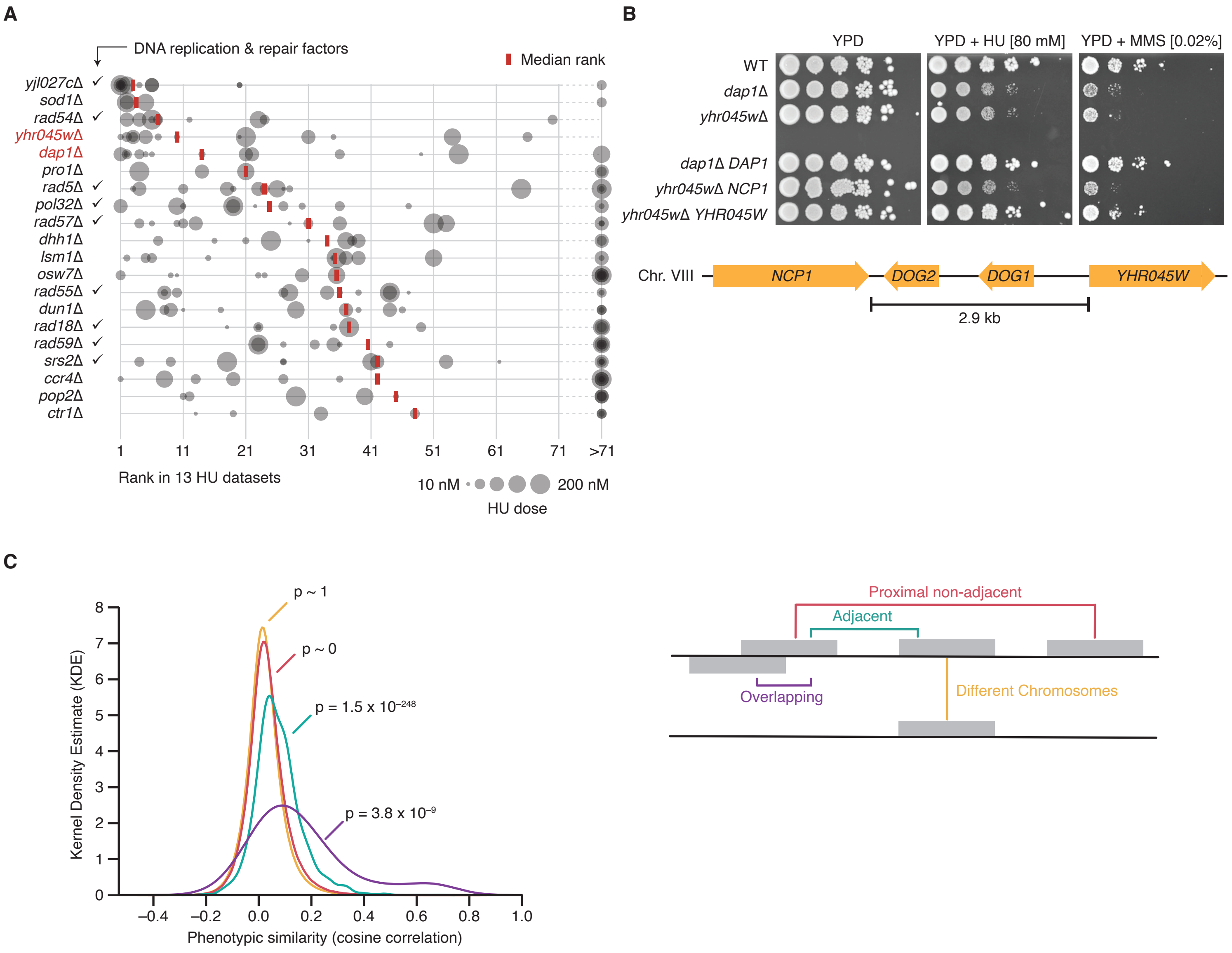


(**A–B**) Evidence from Yeast Phenome and validation experiments suggests that *YHR045W* works with *DAP1* in regulating ergosterol biosynthesis and DNA damage response. (**A**) *yhr045w*Δ and *dap1*Δ are among the top 5 mutants with the highest degree of sensitivity to hydroxyurea (HU). We examined 13 Yeast Phenome screens that tested growth upon exposure to HU at various doses. Mutants were ranked based on their normalized phenotypic values (NPVs) in each screen. The top 20 mutants with the lowest median rank (red line) are shown in the plot. Mutant rank in all 13 datasets is also shown (grey circles). The size of the circles is proportional to the HU dose. *yhr045w*Δ and *dap1*Δ are highlighted in red. (**B**) We experimentally confirmed the sensitivity of *dap1*Δ and *yhr045w*Δ to HU and MMS, and verified that the sensitivity is rescued by the expression of plasmid-borne *DAP1* and *YHR045W*, respectively (Materials & Methods). The sensitivity of *yhr045w*Δ cannot be rescued by the expression of *NCP1*, a gene located 2.9 kb upstream of *YHR045W* and involved in ergosterol biosynthesis like *DAP1*. The inability of *NCP1* to complement *yhr045w*Δ phenotypes indicates that these phenotypes are not caused by a neighboring gene effect. (**C**) The phenotypic similarities of overlapping, immediately adjacent and proximal non-adjacent gene pairs are significantly higher than expected by random chance. A schematic representation of different classes of gene pairs is shown on the right. The distributions of phenotypic similarities for overlapping gene pairs (purple), adjacent gene pairs (cyan), proximal non-adjacent gene pairs (red) and gene pairs located on different chromosomes (yellow) were compared to the overall distribution of all phenotypic similarities (excluding overlapping gene pairs). The p-values for the corresponding 2-sample Kolmogorov-Smirnov tests are reported.

**Figure S9**

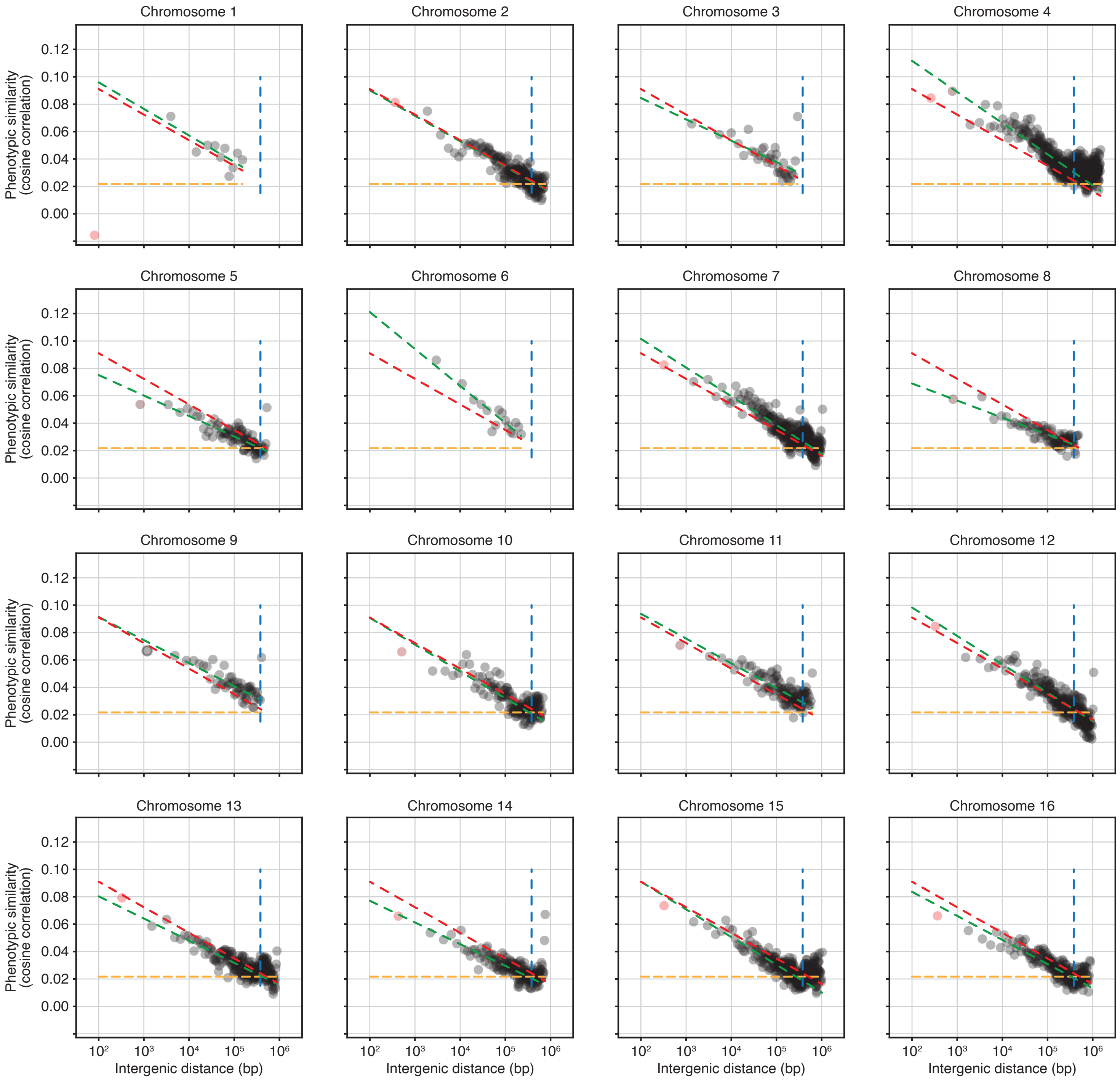


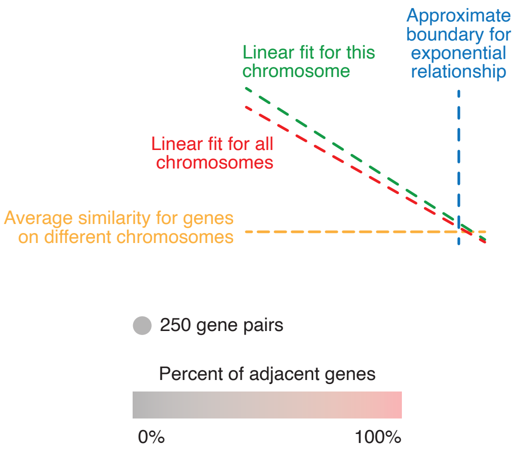


Phenotypic similarity has a strong exponential relationship with chromosomal proximity. This relationship is consistent across all chromosomes examined independently. Gene pairs located on each chromosome were sorted by their intergenic distance and subdivided into groups of 250 pairs. In each group, the average intergenic distance and average phenotypic similarity were computed and plotted on the x and y-axis, respectively. Distance was plotted on a $\log_{10}$ scale. The color of each point indicates the fraction of immediately adjacent genes in the group. The yellow line indicates the average phenotypic similarity for gene pairs located on different chromosomes. The blue line indicates the approximate boundary of the exponential relationship estimated from all chromosomes (380 kb; Fig. 5; Materials & Methods). The green line indicates the linear fit between log10 intergenic distance and phenotypic similarity for all points within the estimated distance boundary (left of the blue line) on each chromosome. As a reference, the red line indicates the same linear fit estimated from all chromosomes (Fig. 5).

**Figure S10**

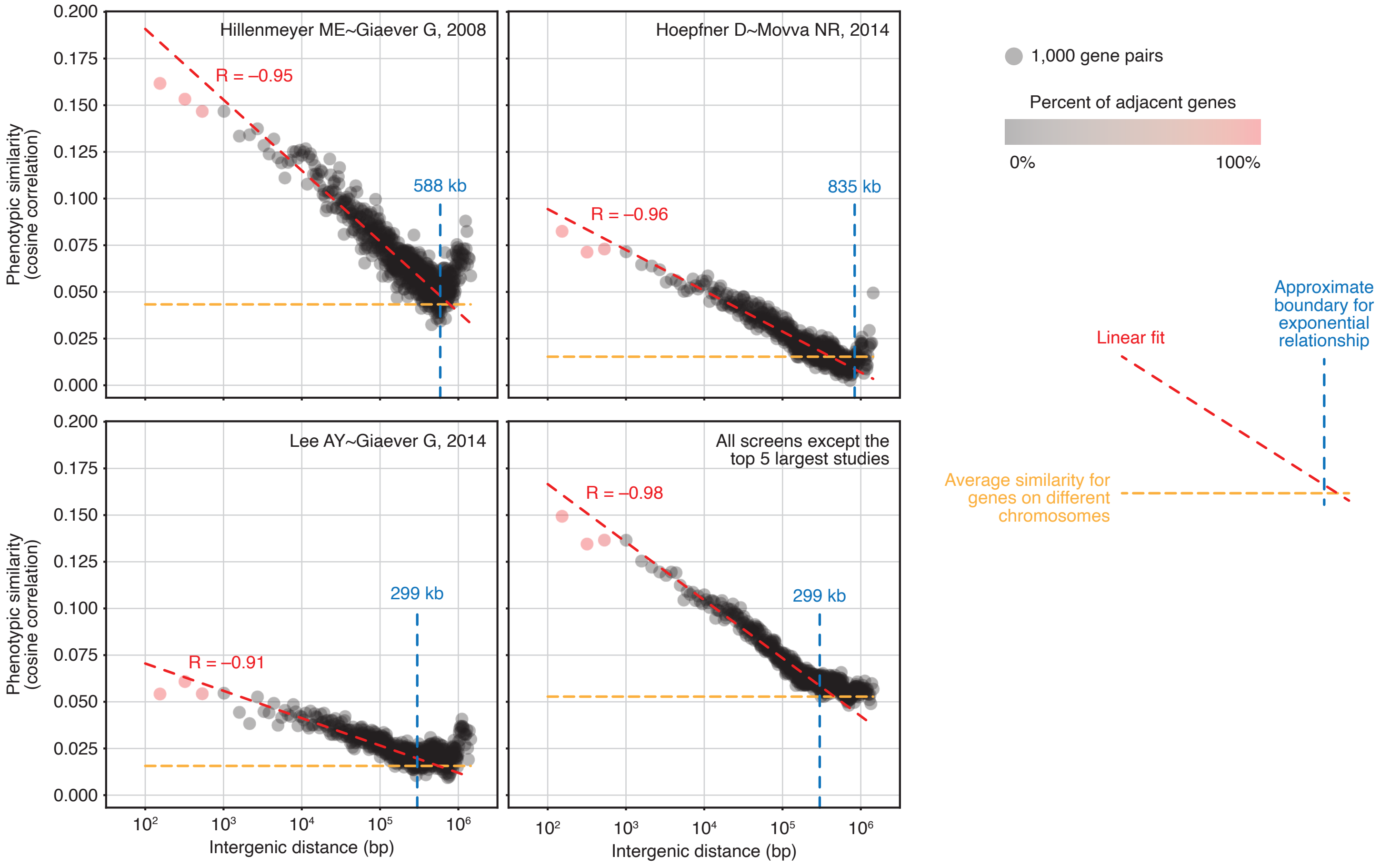


Phenotypic similarity has a strong exponential relationship with chromosomal proximity. This relationship is consistent across several independent subsets of Yeast Phenome data. Phenotypic similarity was computed using only screens from 3 large studies (*6–8*), as well as all screens excluding the top 5 largest chemo-genomics datasets (*6–10*). Gene pairs located on the same chromosome were sorted by their intergenic distance and subdivided into groups of 1,000 pairs. In each group, the average intergenic distance and average phenotypic similarity were computed and plotted on the x and y-axis, respectively. Distance was plotted on a $\log_{10}$ scale. The color of each point indicates the fraction of immediately adjacent genes in the group. The yellow line indicates the average phenotypic similarity for gene pairs located on different chromosomes. The blue line indicates the approximate boundary of the exponential relationship estimated from each subset (Materials & Methods). The red line indicates the linear fit between $\log_{10}$ intergenic distance and phenotypic similarity for all points within the estimated distance boundary (left of the blue line).

### Figure S11

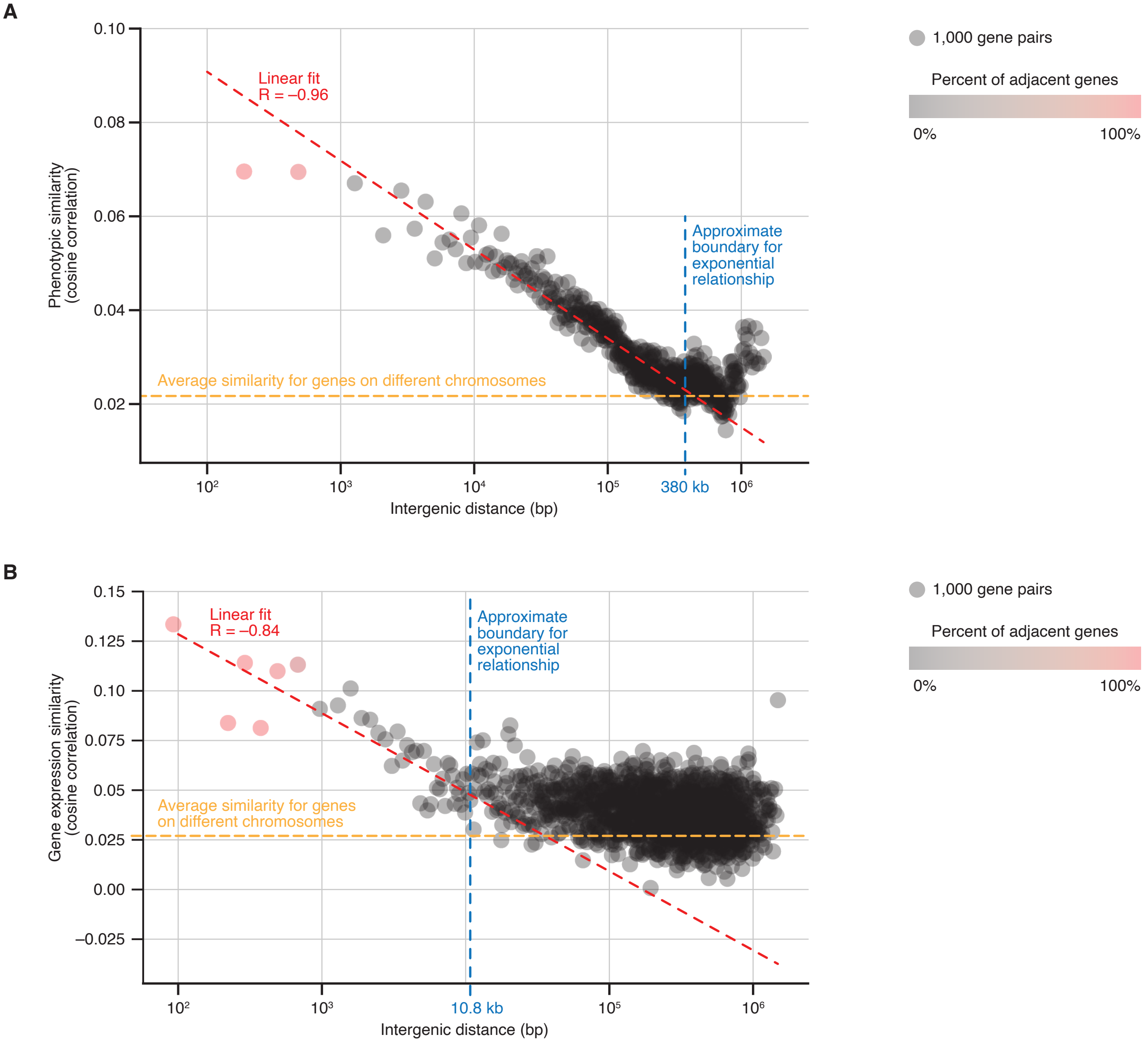


(**A**) Phenotypic similarity has a strong exponential relationship with chromosomal proximity. This relationship persists if we exclude all gene pairs with prior evidence of functional co-clustering (Materials & Methods): members of the same protein complex, metabolic pathway, moderately specific GO biological process term; gene co-expressed or co-regulated by the same transcription factor; paralogous gene pairs. Gene pairs located on the same chromosome were sorted by their intergenic distance and subdivided into groups of 1,000 pairs. In each group, the average intergenic distance and average phenotypic similarity were computed and plotted on the x and y-axis, respectively. Distance was plotted on a $\log_{10}$ scale. The color of each point indicates the fraction of immediately adjacent genes in the group. The yellow line indicates the average phenotypic similarity for gene pairs located on different chromosomes. The blue line indicates the approximate boundary of the exponential relationship estimated from each subset (Materials & Methods). The red line indicates the linear fit between $\log_{10}$ intergenic distance and phenotypic similarity for all points within the estimated distance boundary (left of the blue line). (**B**) Similarity of gene expression across multiple experimental conditions (*41*) is also related to chromosomal proximity. However, this relationship has a much shorter range (10.8 kb) than phenotypic similarity (380 kb; Materials & Methods). Gene pairs located on the same chromosome were sorted by their intergenic distance and subdivided into groups of 1,000 pairs. In each group, the average intergenic distance and average gene co-expression (measured by cosine similarity) were computed and plotted on the x and y-axis, respectively. Distance was plotted on a $\log_{10}$ scale. The color of each point indicates the fraction of immediately adjacent genes in the group. The yellow line indicates the average phenotypic similarity for gene pairs located on different chromosomes. The blue line indicates the approximate boundary of the exponential relationship estimated from each subset (Materials & Methods). The red line indicates the linear fit between $\log_{10}$ intergenic distance and phenotypic similarity for all points within the estimated distance boundary (left of the blue line).

*Table S1 – List of Yeast Phenome screens for respiratory metabolism*

| Dataset ID | Paper | Collection | Growth assay | Medium | Data type | Notes |
|---|---|---|---|---|---|---|
| 470 | Dimmer KS~Westermann B, 2002 | Hom | Undefined | YPG 2% | Discrete | |
| 4837 | Steinmetz LM~Davis RW, 2002 | Hom | Pooled culture | YPG 3% | Quantitative | |
| 5000 | Dudley AM~Church GM, 2005 | Hom | Spot assay | YPG 3% | Discrete | |
| 417 | Luban C~Schmidt U, 2005 | Hap a | Undefined | YPG 3% | Discrete | |
| 21955 | Kuepfer L~Blank LM, 2005 | Hap a | Undefined | MM (Verduyn mix) + His + Leu + Met + Ura, Glycerol (2%) | Discrete | Excluded from analysis (SC medium) |
| 1050, 1051 | Hillenmeyer ME~Giaever G, 2008 | Hom | Pooled culture | YPG | Quantitative | Excluded from analysis (unknown YPG dose) |
| 158 | Merz S~Westermann B, 2009 | Hap alpha | Spot assay | YPG 3% | Discrete | |
| 16489 | Qian W~Zhang J, 2012 | Hom | Pooled culture | YPG 5% | Quantitative | |

| | | | | | | |
|---|---|---|---|---|---|---|
| 16369, 16388 | Galardini M~Beltrao P, 2019 | Hap a | Colony size | SC + G 2% | Quantitative | Excluded from analysis (SC medium) |
| 22018 | Acton E~Giaever G, 2017 | Hom | Pooled culture | YPG 3% | Quantitative | |
| 21874 | Stenger M~Westermann B, 2020 | Hap a | Colony size | YPG 3% | Discrete | |

Only experiments that employed YP-based media and glycerol as the sole carbon source were included in this analysis. Yeast Phenome contains additional data for growth on synthetic media (partial and complete) and media supplemented with other carbon sources (e.g., glucose, ethanol).

***Table S2 – Analysis of phenotype-function consistency for knock-out mutants carrying secondary mutations***

| | | **Phenotype-function consistency** | | | |
|---|---|---|---|---|---|
| | | **Yes** | **No** | **Total** | **Risk** |
| **Secondary mutations** | **Yes** | A = 72<br>$A_0$ = 66 | B = 31<br>$B_0$ = 37 | 103 | ARM = b / (a+b) = 0.3 |
| | **No** | C = 60<br>$C_0$ = 66 | D = 44<br>$D_0$ = 38 | 104 | ARW = d / (c+d) = 0.42 |
| | **Total** | 132 | 75 | 207 | |

The relative risk of a secondary mutation affecting the phenotypes is given by RR = ARM / ARW = 0.3 / 0.42 = 0.711 (values below 1 indicate that secondary mutations help, do not harm, phenotypic profiles).

To estimate the confidence intervals around this relative risk, we can use the Taylor series approximate variance (*94*). The two-sided 95% confidence limits are given by:

$$CI = e^{lnRR \pm 1.96\sqrt{\frac{1-ARM}{b}+\frac{1-ARW}{d}}}$$

So, in this case, RR = 0.711, CI 95%: [0.491, 1.030]. That means that, with 95% confidence, the relative risk of a secondary mutation to negatively impact the phenotypic profile of a knock-out mutant is at most 3%.

***Table S3 – List of uncharacterized ORFs with at least 1% pleiotropy and strong profile similarity to verified ORFs***

| | Top correlated gene | | | | |
|---|---|---|---|---|---|
| **Uncharacterized ORF** | **ORF** | **Gene name** | **Correlation (mean)** | **Correlation (std. dev.)** | **Phenotype rate (%)** |
| *YLR261C* | *YLR262C* | *YPT6* | 0.689 | 0.05 | 0.13 |
| *YBR062C* | *YGL131C* | *SNT2* | 0.667 | 0.034 | 0.01 |
| *YGL117W* | *YDR354W* | *TRP4* | 0.66 | 0.031 | 0.04 |
| *YHR045W* | *YPL170W* | *DAP1* | 0.582 | 0.068 | 0.03 |
| *YDR114C* | *YER050C* | *RSM18* | 0.535 | 0.038 | 0.03 |
| *YIL077C* | *YIL041W* | *GVP36* | 0.515 | 0.099 | 0.02 |
| *YIL029C* | *YOR043W* | *WHI2* | 0.489 | 0.065 | 0.02 |
| *YKR073C* | *YKR072C* | *SIS2* | 0.476 | 0.037 | 0.02 |
| *YIL014C-A* | *YIL060W* | *YIL060W* | 0.471 | 0.037 | 0.02 |
| *YFL034W* | *YHL019C* | *APM2* | 0.458 | 0.081 | 0.02 |
| *YNL184C* | *YNL252C* | *MRPL17* | 0.445 | 0.046 | 0.03 |
| *YHL029C* | *YNL056W* | *OCA2* | 0.441 | 0.05 | 0.01 |
| *YCR087C-A* | *YCR045C* | *RRT12* | 0.439 | 0.04 | 0.02 |
| *YJL193W* | *YJL155C* | *FBP26* | 0.437 | 0.073 | 0.02 |
| *YLL030C* | *YIL121W* | *QDR2* | 0.418 | 0.065 | 0.01 |
| *YPL056C* | *YPL057C* | *SUR1* | 0.417 | 0.125 | 0.01 |
| *YCR050C* | *YCR045C* | *RRT12* | 0.417 | 0.047 | 0.03 |
| *YGL088W* | *YDR450W* | *RPS18A* | 0.412 | 0.033 | 0.02 |
| *YLR331C* | *YLR332W* | *MID2* | 0.405 | 0.069 | 0.02 |
| *YOR183W* | *YPR106W* | *ISR1* | 0.396 | 0.035 | 0.03 |
| *YCL001W-A* | *YCR026C* | *NPP1* | 0.395 | 0.045 | 0.03 |

| | | | | | |
|---|---|---|---|---|---|
| *YCR007C* | *YDL010W* | *GRX6* | 0.391 | 0.059 | 0.01 |
| *YNL140C* | *YHR167W* | *THP2* | 0.387 | 0.057 | 0.03 |
| *YPR089W* | *YDL226C* | *GCS1* | 0.364 | 0.06 | 0.01 |
| *YBR027C* | *YDL066W* | *IDP1* | 0.362 | 0.049 | 0.01 |
| *YCR085W* | *YBR295W* | *PCA1* | 0.356 | 0.045 | 0.01 |
| *YEL033W* | *YIR026C* | *YVH1* | 0.352 | 0.035 | 0.03 |
| *YKL121W* | *YBR208C* | *DUR1,2* | 0.347 | 0.051 | 0.01 |
| *YER077C* | *YNL177C* | *MRPL22* | 0.346 | 0.039 | 0.04 |
| *YGL007W* | *YMR216C* | *SKY1* | 0.341 | 0.041 | 0.05 |
| *YBR284W* | *YCR045C* | *RRT12* | 0.336 | 0.047 | 0.02 |
| *YEL028W* | *YBR149W* | *ARA1* | 0.33 | 0.064 | 0.01 |
| *YCR025C* | *YCR045C* | *RRT12* | 0.329 | 0.042 | 0.01 |
| *YLR426W* | *YDL066W* | *IDP1* | 0.327 | 0.035 | 0.04 |
| *YER084W* | *YDL100C* | *GET3* | 0.326 | 0.047 | 0.01 |
| *YMR262W* | *YDR458C* | *HEH2* | 0.322 | 0.047 | 0.01 |
| *YML037C* | *YPR029C* | *APL4* | 0.319 | 0.044 | 0.01 |
| *YCR051W* | *YBR278W* | *DPB3* | 0.314 | 0.037 | 0.01 |
| *YLR358C* | *YJL080C* | *SCP160* | 0.31 | 0.046 | 0.06 |
| *YML122C* | *YLR372W* | *ELO3* | 0.304 | 0.041 | 0.12 |
| *YLR407W* | *YJR083C* | *ACF4* | 0.299 | 0.062 | 0.01 |
| *YDR525W* | *YOR085W* | *OST3* | 0.296 | 0.044 | 0.02 |
| *YHL044W* | *YJL192C* | *SOP4* | 0.29 | 0.04 | 0.01 |
| *YIL067C* | *YCR009C* | *RVS161* | 0.238 | 0.045 | 0.02 |
| *YHL005C* | *YMR256C* | *COX7* | 0.236 | 0.042 | 0.01 |
| *YLR125W* | *YDL184C* | *RPL41A* | 0.176 | 0.032 | 0.01 |

*Table S4 – List of yeast strains used for validation experiments*

| Strain ID | Genotype |
|---|---|
| Yeast knock-out collection (YKO) | MATa *orf*Δ::KanMX *his3*Δ1 *leu2*Δ0 *met15*Δ0 *ura3*Δ0 |
| Prototrophic deletion collection (PDC) | MATa *orf*Δ::KanMX *can1*Δ::*STE2pr*-Sp_*his5* *his3*Δ1 *lyp1*Δ0 |
| ABY001 | PDC hoΔ::KanMX |
| ABY002 | PDC *dap1*Δ::KanMX |
| ABY003 | PDC *yhr045w*Δ::KanMX |
| ABY004 | PDC *ygl117w*Δ::KanMX |
| ABY005 | PDC *dap1*Δ::KanMX *yhr045w*Δ::NatMX |
| ABY006 | ABY001 [MoBY-2μ-*LEU2*-*ERG11*] |
| ABY007 | ABY002 [MoBY-2μ-*LEU2*-*ERG11*] |
| ABY008 | ABY003 [MoBY-2μ-*LEU2*-*ERG11*] |
| ABY009 | ABY002 [pRS412-NatNT2-*DAP1*] |
| ABY010 | ABY003 [pRS412-NatNT2-*YHR045W*] |
| ABY011 | ABY003 [pRS412-NatNT2-*NCP1*] |
| ABY012 | ABY004 [MoBY-2μ-*LEU2*-*YGL117W*] |
